# A Protein Engineering Approach for Uncovering Cryptic Ubiquitin-binding Sites: from a Ubiquitin-Variant Inhibitor of APC/C to K48 Chain Binding

**DOI:** 10.1101/669846

**Authors:** Edmond R. Watson, Christy R. R. Grace, Wei Zhang, Darcie J. Miller, Iain F. Davidson, J. Rajan Prabu, Shanshan Yu, Derek L. Bolhuis, Elizaveta T. Kulko, Ronnald Vollrath, David Haselbach, Holger Stark, Jan-Michael Peters, Nicholas G. Brown, Sachdev S. Sidhu, Brenda A. Schulman

## Abstract

Ubiquitin-mediated proteolysis is a fundamental mechanism used by eukaryotic cells to maintain homeostasis and protein quality, and to control timing in biological processes. Two essential aspects of ubiquitin regulation are conjugation through E1-E2-E3 enzymatic cascades, and recognition by ubiquitin-binding domains. An emerging theme in the ubiquitin field is that these two properties are often amalgamated in conjugation enzymes. In addition to covalent thioester linkage to ubiquitin’s C-terminus for ubiquitin transfer reactions, conjugation enzymes often bind non-covalently and weakly to ubiquitin at “exosites”. However, identification of such sites is typically empirical and particularly challenging in large molecular machines. Here, studying the 1.2 MDa E3 ligase Anaphase-Promoting Complex/Cyclosome (APC/C), which controls cell division and many aspects of neurobiology, we discover a method for identifying unexpected ubiquitin-binding sites. Using a panel of ubiquitin variants (UbVs) we identify a protein-based inhibitor that blocks ubiquitin ligation to APC/C substrates in vitro and ex vivo. Biochemistry, NMR, and cryo EM structurally define the UbV interaction, explain its inhibitory activity through binding the surface on the APC2 subunit that recruits the E2 enzyme UBE2C, and ultimately reveal that this APC2 surface is also a ubiquitin-binding exosite with preference for K48-linked chains. The results provide a new tool for probing APC/C activity, have implications for the coordination of K48-linked Ub chain binding by APC/C with the multistep process of substrate polyubiquitylation, and demonstrate the power of UbV technology for identifying cryptic ubiquitin binding sites within large multiprotein complexes.

**SIGNIFICANCE STATEMENT:** Ubiquitin-mediated interactions influence numerous biological processes. These are often transient or a part of multivalent interactions. Therefore, unmasking these interactions remains a significant challenge for large, complicated enzymes such as the Anaphase-Promoting Complex/Cyclosome (APC/C), a multisubunit RING E3 ubiquitin (Ub) ligase. APC/C activity regulates numerous facets of biology by targeting key regulatory proteins for Ub-mediated degradation. Using a series of Ub variants (UbVs), we identified a new Ub-binding site on the APC/C that preferentially binds to K48-linked Ub chains. More broadly, we demonstrate a workflow that can be exploited to uncover Ub-binding sites within ubiquitylation machinery and other associated regulatory proteins to interrogate the complexity of the Ub code in biology.

## INTRODUCTION

Post-translational modification by ubiquitin (Ub) regulates numerous eukaryotic cellular processes including cell division, signal transduction, transcription, translation, protein trafficking, and more. The exceptional regulatory potential of the Ub system stems from both a vast enzymatic system that links Ub to specific protein targets, and enormous diversity in forms of ubiquitylation. Ub is not a singular modification, but is often itself modified by additional Ubs in the form of “chains”, where Ubs are linked via one of eight different primary amino groups (lysine side-chains or the N-terminus) on another Ub. Precise ubiquitylation is catalyzed by specific combinations of E3 Ub ligases and E2 Ub conjugating enzymes, and the resultant Ub marks are recognized by Ub-binding domains that impart distinct fates to modified proteins (1–8). As examples, K11- and K48-linked Ub chains – particularly in combination with each other – elicit proteasomal degradation, whereas K63-linked chains often modulate subcellular location or assembly and disassembly of macromolecular complexes.

E3 enzymes achieve ubiquitylation by recruiting one or more sequence motifs, termed “degrons”, in a substrate, and promoting Ub transfer through one of various mechanisms determined by the type of catalytic domain. E3s harboring “HECT” and “RBR” catalytic domains promote ubiquitylation through 2-step reactions involving formation of a thioester-linked intermediate between the E3 and Ub’s C-terminus: first, Ub is transferred from an E2~Ub intermediate (“~” refers to thioester bond) to the E3 catalytic Cys; Ub is subsequently transferred from the E3’s Cys to the substrate (9–13). Alternatively, the majority of E3s, including an estimated ≈500 RING E3s in humans, do not directly relay Ub themselves, but instead activate Ub transfer from the catalytic Cys of a Ub-carrying enzyme, which is typically an E2, but can also be a thioester-forming E3 (14, 15). Notably, many RING E3s are multifunctional, interacting with several distinct Ub carrying enzymes to link various forms of Ub to myriad substrates. In many cases, a substrate undergoes polyubiquitylation by different combinations of enzymes acting in series. For example, some RING E3s employ different E2s with one first priming a substrate directly with Ub, and then the other extending Ub chains from substrate-linked Ubs (16, 17).

Recent studies have shown that the catalytic domains of several E3 and E2 enzymes not only carry Ub covalently at their active sites to mediate ubiquitylation, but also bind Ub noncovalently at “exosites” remote from their hallmark catalytic surfaces (18). Noncovalent Ub binding can influence targeting, processivity, and rates of ubiquitylation reactions. As examples, noncovalent Ub binding to the E2 UBE2D2 allosterically modulates binding to partner RING domain E3s (19, 20), while noncovalent binding to HECT E3s in the NEDD4-family can modulate catalytic activity and/or processivity of substrate ubiquitylation (21). The first RING domain discovered to possess a Ub-binding exosite was APC11, a subunit of one of the most complicated and unusual E3 ligases, the Anaphase-Promoting Complex/Cyclosome (APC/C) (22). APC/C, a 1.2 MDa complex comprised of 19 core polypeptides including APC11 and the cullin-like APC2, controls cell division by promoting ubiquitin-dependent turnover of cyclins and other key cell cycle regulators (reviewed in (23, 24)). Recent structural studies have revealed how APC/C is activated through binding to a coactivator (either CDC20 or CDH1), which recruits substrate degrons and conformationally activates the APC2-APC11 cullin-RING catalytic core (25); and how the APC2-APC11 catalytic core, in turn, differentially recruits and activates distinctive E2~Ub intermediates to generate various ubiquitylated products (26–28). Human APC/C employs the E2 UBE2C to prime substrates with one or multiple individual Ubs or short polyUb chains, and the E2 UBE2S to extend chains with K11 linkages (29–32). When UBE2S encounters substrates modified with K48-linked Ub chains generated by UBE2C, “branched” chains containing K48 and K11 linkages are produced, which are particularly potent at directing proteasomal degradation (33).

APC11’s Ub-binding exosite makes distinct contributions to the reactions with the different E2s (Fig. 1a). With UBE2C, in a process called “Processive Affinity Amplification”, this exosite provides additional affinity for Ub-linked substrates, which increases propensity for further modification (28, 34). In a different, crucial role in APC/C-dependent polyubiquitylation with UBE2S, APC11’s Ub-binding exosite recruits substrate-linked Ubs for K11-linked Ub chain elongation (22, 35). In addition to the key roles played by Ub binding to APC11, single molecule studies indicated the presence of other, unidentified, Ub-binding sites on APC/C (28).

**Figure 1.**
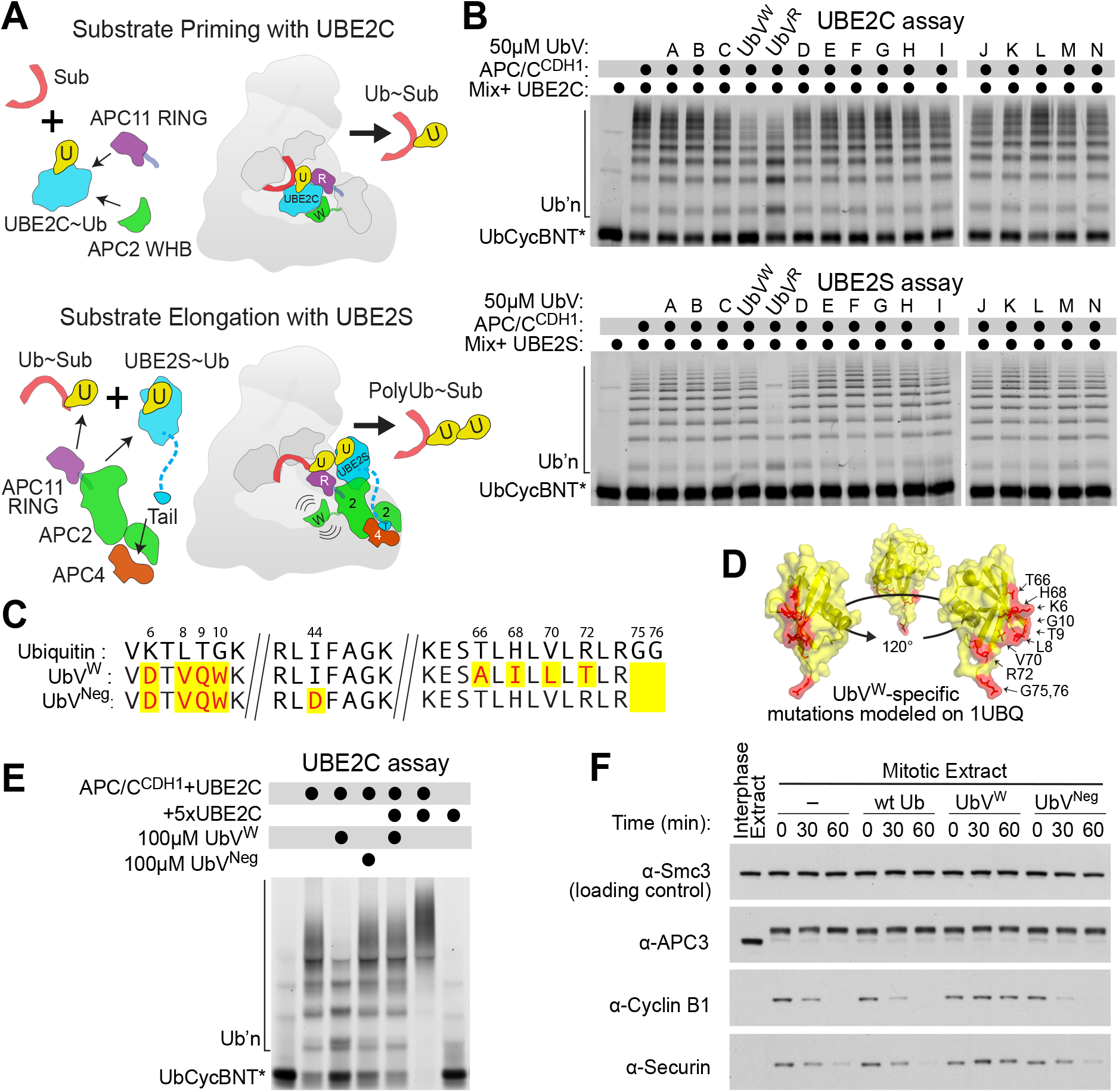
Discovery of a Ubiquitin Variant inhibitor (UbV^W^) of APC/C activity with UBE2C. (A) Cartoon illustrating the distinct APC/C ubiquitylation mechanisms. In a substrate priming reaction (top), UBE2C (cyan) is harnessed by APC2 WHB (green) and the canonical APC11 RING residues (purple) to catalyze transfer of Ubiquitin (yellow) to substrate (red). In a Ub chain elongation reaction (bottom), UBE2S is harnessed at a distal site and utilizes distinct surfaces of APC2 (green), APC4 (orange), and APC11 (purple) to generate polyubiquitin chains on existing Ub conjugates (yellow). (B) A panel of Ubiquitin Variant (UbV) proteins was assayed for effects on ubiquitylation using UbCycBNT* (Ub-fusion to fluorescent CyclinB N-terminal domain), an APC/C^CDH1^ substrate suitable for modification by both the substrate-priming E2 UBE2C (top) and the Ub chain elongating E2 UBE2S (bottom). Ubiquitylated products are separated via 15% SDS-PAGE and visualized in a Typhoon biomolecular fluorescence imager. (C) Sequence alignment of Ubiquitin, UbV^W^ and negative control mutant UbV^Neg^, with divergent residues highlighted in yellow. (D) Sequence divergence in UbV^W^ are shown in red, mapped on the structure of ubiquitin (PDB: 1UBQ). (E) Ubiquitylation of UbCycBNT* (Ub-fusion to fluorescent CyclinB N-terminal domain) is monitored in the presence of excess UBE2C, UbV^W^, or a negative control mutant UbV^Neg^. UbV^W^ inhibits in the presence of excess UBE2C suggesting indirect inhibition. (F) Effects of UbV^W^, or ubiquitin or UbV^Neg^, on degradation of exogenous APC/C substrates Cyclin B1 and Securin in extracts from mitotic *Xenopus* eggs, monitored by western blot. Antibody specificities are denoted to the left, with SMC3 as loading control, and the slower migrating, phosphorylated APC3 indicating active APC/C in the extracts.

Although it is of great interest to identify Ub-binding domains, it can be challenging to find these interactions de novo. One obstacle is that the affinities of interactions between Ub and binding domains are typically extremely low, with Kds often in the millimolar range (36). Such low affinities generally reflect a single Ub being only one of many elements contributing to avid, multisite interactions (37). As a consequence, many Ub-binding sites have been identified by yeast 2-hybrid, pulldowns, or NMR experiments with high concentrations of interacting domains. However, in the absence of sequence motifs indicating the presence of a Ub-binding domain, it is an arduous task to find such interactions within massive assemblies that cannot be produced in large quantities at high concentrations for techniques such as NMR. Indeed, we previously discovered the Ub-binding exosite fortuitously, through alanine-scanning mutagenesis of APC11’s RING domain and assaying enzymatic activity with UBE2S in the context of the fully assembled, recombinant >1.2 MDa APC/C^CDH1^ complex (22).

We considered that it might be possible to use “Ub variant” (UbV) technology to identify a new Ub-binding site within APC/C. Unlike Ub, which is constrained in sequence and structure by the requirement to bind a massive number of partner proteins, sequence variants have the potential to make distinctive contacts that increase affinity for particular partners. We and others previously performed phage display to select variants binding to a variety of E3 ligases and deubiquitylating enzymes (DUBs), and previously identified Ub-binding domains, to test effects of perturbing Ub interactions with known interaction sites (21, 28, 38–44). Indeed, our UbV^R^, selected from among 10^10^ Ub variants (UbVs) displayed on phage as binding APC11’s RING domain, proved useful for uncovering the contribution of APC11’s Ub-binding exosite to processive affinity amplification of activity with UBE2C, and enabled obtaining cryo EM data visualizing UBE2S bound to the APC2-APC11 catalytic core of APC/C (28).

Here, we utilize UbV technology with a different goal: we tested for perturbation of E3 activity without prior knowledge of a specific Ub-binding site. In so doing, we generated an inhibitor of substrate priming by UBE2C that stabilizes APC/C targets in mitotic extracts. Remarkably, this strategy also identified the UBE2C-binding site on the APC2 cullin domain as a previously unknown Ub-binding exosite with preference for K48-linked chains. The results have implications for the interplay between polyubiquitylated substrates, E2s, and other factors regulating APC/C activity through interactions with the cullin subunit. Furthermore, we provide a route for identifying novel Ub-binding sites within massive multiprotein assemblies.

## RESULTS

### Identification of a Ubiquitin variant (UbV) inhibiting ubiquitin-mediated proteolysis of APC/C-UBE2C substrates

During the course of characterizing effects of UbV^R^, which binds APC11’s RING domain with ≈1 micromolar affinity, we monitored APC/C^CDH1^-catalyzed ubiquitylation in the presence of excess individual UbVs from a panel of purified proteins available in the laboratory. UBE2C-dependent substrate priming and UBE2S-dependent Ub chain extending activities (Fig. 1a) were assayed in parallel using the model substrate “UbCycBNT*”, consisting of Ub fused upstream of the N-terminal domain of Cyclin B, which contains a D-box “degron”, and a C-terminal fluorophore. In the control reactions recapitulating our prior results (28), UbV^R^ decreased processivity with UBE2C, as shown by conversion of substantial substrate to products with fewer ubiquitins (Fig. 1b). In addition, all activity with UBE2S is impaired by UbV^R^ competing with the acceptor Ub for binding to the APC11 RING domain. Most UbVs had little effect on either reaction, which was not surprising given that none of them were selected specifically for binding to domains from APC/C^CDH1^. Serendipitously, however, one UbV (termed UbV^W^, for reasons described below) displayed a distinctive profile: selective inhibition of reactions with UBE2C with no obvious effect on polyubiquitylation with UBE2S (Fig. 1b). Relative to wt Ub, UbV^W^ displays mutations primarily in the “Leu8 loop” between the two N-terminal β-strands, and in the C-terminal tail, and retains the I44 hydrophobic patch known to mediate most interactions, albeit with L8V, H68I, and V70L substitutions (Fig. 1c,1d).

Consistent with the notion that UbV^W^ specifically antagonizes APC/C with UBE2C, reactions with relatively higher concentrations of UBE2C showed increased ubiquitylation, which were attenuated by the presence of UbV^W^ (Fig. 1e). An additional UbV selected as a negative control (“UbV^Neg^”) showed no effect on these reactions, and excess UBE2C alone does not catalyze ubiquitylation in the absence of APC/C.

To evaluate APC/C and UBE2C activity in a more physiological setting, we examined effects of UbV^W^ on degradation of well-characterized substrates, Cyclin B1 and Securin, in mitotic *Xenopus* egg extracts. These extracts were previously shown to retain key cellular requirements for APC/C-dependent degradation during the cell cycle, including substrate priming activity from collaboration with endogenous UBE2C present in the extracts (29, 45, 46). Importantly, addition of UbV^W^, but not UbV^Neg^, stabilized the cell cycle proteins from APC/C-UBE2C-dependent degradation (Fig. 1f).

### UbV^W^ hijacks APC2’s WHB domain to compete with binding to UBE2C

To gain mechanistic insights into how UbV^W^ could selectively inhibit APC/C activity with UBE2C, we performed kinetic analyses. Adding saturating concentrations of UbV^W^ while titrating UBE2C caused a 5-fold increase in *K*_*m*_ with only a minor effect on the *V*_*max*_ (Fig. 2a), implying that UbV^W^ antagonizes UBE2C binding to APC/C.

**Figure 2.**
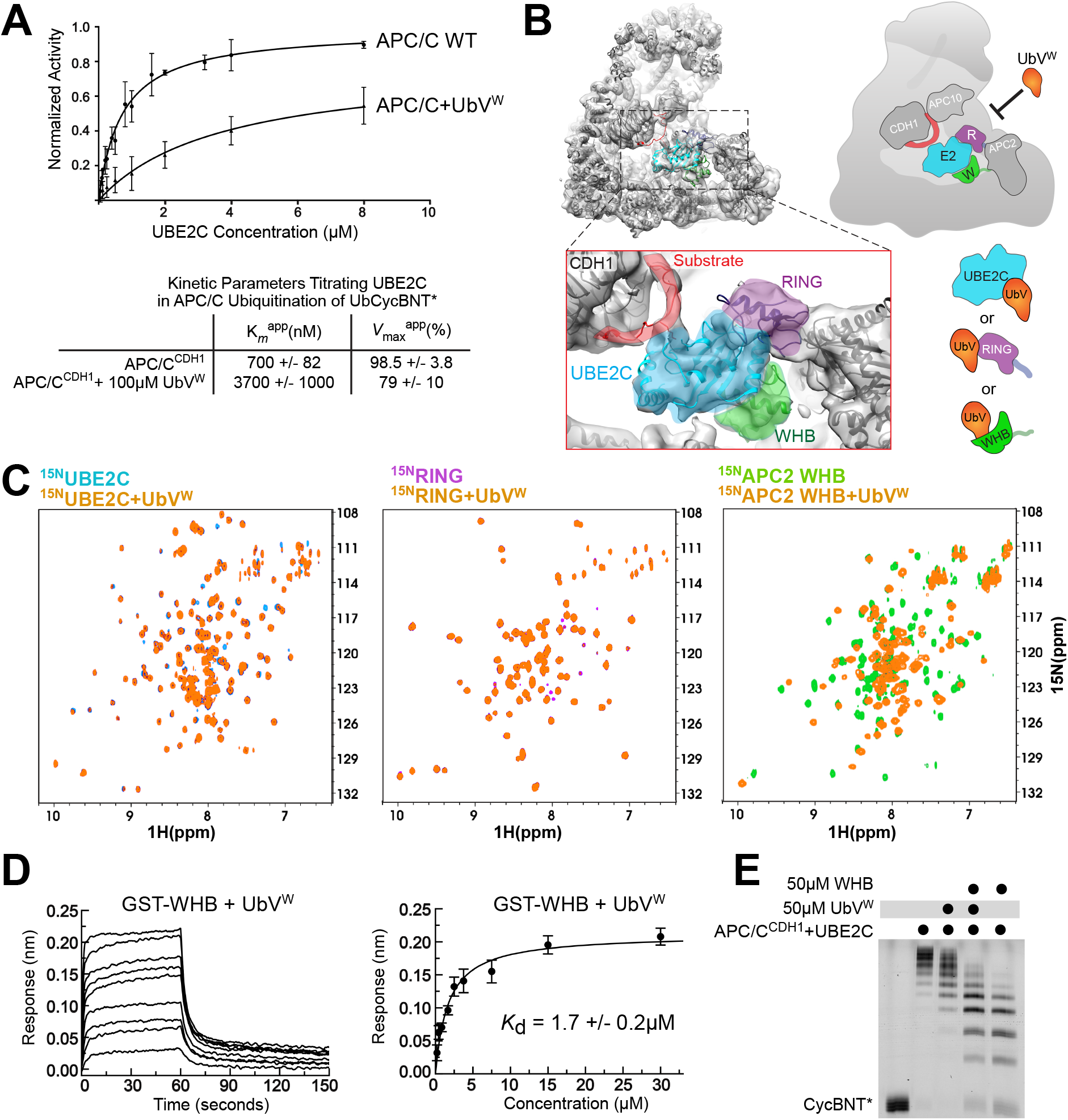
UbV^W^ directly interacts with APC/C to antagonize UBE2C activity. (A) Curve fits (top) and kinetic parameters (bottom) testing effects of saturating UbV^W^ on titrations of UBE2C in APC/C^CDH1^-mediated ubiquitylation of UbCycBNT*. SEM, n ≥ 3. (B) Potential mechanisms of UbV^W^ inhibition of APC/C activity with UBE2C, based on prior EM map [EMDB:3432] of UBE2C bound to APC/C^CDH1^-substrate, with closeup of substrate (red), UBE2C (cyan), APC11 RING (purple) and APC2 WHB domain (green). Right, cartoons illustrating three possible mechanisms for UbV inhibition of UBE2C recruitment by associating with UBE2C directly, or with APC11 RING domain, or with APC2 WHB domain. (C) Left, overlaid [^15^N, ^1^H] TROSY spectra of ^15^N-labeled UBE2C alone (100μM, cyan) or with UbV^W^ (1:2, orange). Middle, overlaid [^15^N, ^1^H] TROSY spectra of ^15^N-labeled APC11 RING alone (100μM, purple) or with UbV^W^ (1:2, orange). Right, overlaid [^15^N, ^1^H] HSQC spectra of ^15^N-labeled APC2 WHB alone (100μM, green) or with UbV^W^ (1:2, orange). (D) Representative sensorgram and curve fit for binding measured by Bio-Layer Interferometry (BLI) with UbV^W^ titrated in solution versus immobilized GST-APC2 WHB domain. SEM, n ≥ 2. (E) Ubiquitylation of CycBNT* (Fluorescent CyclinB N-terminal domain) by APC/C and UBE2C is monitored in the presence of UbV^W^, excess isolated WHB domain, or a combination of both inhibitors. Alleviation of inhibition when UbV^W^ and isolated WHB are used in combination suggests direct binding.

Three potential explanations for this result are suggested from prior structures of UBE2C bound to APC/C (Fig. 2b) (26, 27). First, UbV^W^ could bind directly to UBE2C to compete with APC/C binding. Second, UbV^W^ could block the canonical E2-binding site on the APC11 RING domain. Third, UbV^W^ could prevent the distinctive recruitment of UBE2C that is mediated by the so-called “WHB domain” from APC2. To distinguish between these possibilities, we performed NMR experiments examining the effects on ^15^N-^1^H TROSY or HSQC spectra when UbV^W^ was added to ^15^N-labeled versions of UBE2C, APC11 RING domain, or APC2 WHB domain. Only the latter spectrum showed substantial chemical shift perturbations, indicating that UbV^W^ binds directly to the WHB domain from APC2 (Fig. 2c). Indeed, a *K*_d_ of ~1.7μM was measured by Bio-Layer Interferometry upon titrating UbV^W^ with immobilized GST-APC2 WHB (Fig. 2d). Hence, the name UbV^W^ reflects to binding to APC2’s **<u>W</u>**HB domain. Interestingly, the APC2 WHB was already known to be a multifunctional protein interaction domain. In addition to its role in recruiting UBE2C to catalyze substrate ubiquitylation, this domain binds the BUBR1 subunit of Mitotic Checkpoint Complex during APC/C inhibition from the spindle assembly checkpoint (47, 48).

An in vitro ubiquitylation assay with opposing inhibitors was used to further test if UbV^W^ competing with UBE2C for APC2’s WHB domain is responsible for the inhibition of APC/C activity. UBE2C-dependent ubiquitylation of a substrate (fluorescent Cyclin B N-terminal domain, “CycBNT*) is inhibited both by UbV^W^, which presumably binds the APC2 WHB domain in context of the APC/C^CDH1^ complex, and by the isolated APC2 WHB domain, which binds UBE2C and prevents its binding to APC/C (27). However, adding UbV^W^ partially ameliorated the inhibition exerted by the isolated APC2 WHB domain, consistent with UbV^W^ binding in effect liberating UBE2C to perform ubiquitylation (Fig. 2e). Taken together, the results indicate that UbV^W^ binds APC2’s WHB domain within the APC/C^CDH1^ complex in a manner that competes with the binding of UBE2C.

### A domain-swapped dimer structure enables UbV^W^ to bind the APC/C with high affinity

As a first step toward understanding the structural basis for its effects, we determined a crystal structure of UbV^W^ at 2.8 Å resolution (Supplemental Table 1). The structure revealed two striking features. First, the crystal contains an oligomeric crystal packing assembly with eight copies in the asymmetric unit (Fig. 3a). Analytical ultracentrifugation and size exclusion chromatography-multiangle light scattering confirmed a higher-order oligomer (tetramer) in solution (Fig. 3b, Supplemental Fig. 1a). Second, the model revealed that the UbV forms a symmetric domain-swapped dimer where the normal Ub β1-β2-loop comprised of residues 6-9 is extended such that the β1 strand from one subunit forms an antiparallel β-sheet with the opposite subunit (Fig. 3c). In this arrangement, we refer to a “monomer” as the portion of the dimer corresponding to the Ub-like fold and the residues involved in the domain swap. This comprises residues 1-10 of one protomer (A) and 8-74 of the opposite protomer (B) in the domain-swapped dimer, where Val8 of protomer A indirectly stabilizes the fold, and Val8-Trp10 of both protomers form the β-sheet extension mediating the domain swap. The globular portion (residues 1-70) of the “monomer” superimposes on the corresponding portion of Ub (residues 1-70) with 1.2Å backbone RMSD.

**Figure 3.**
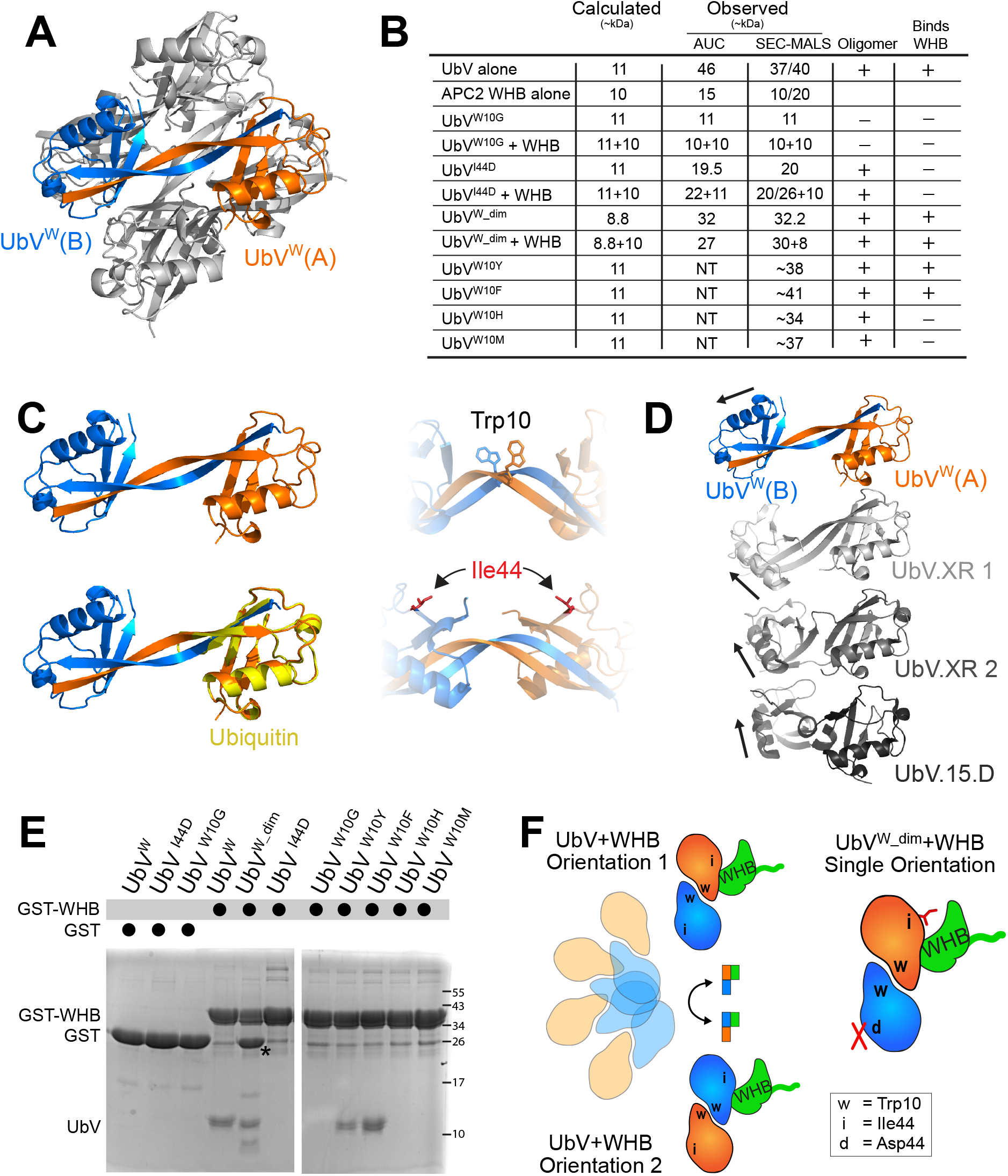
Structure of UbV^W^ reveals dynamic association with APC/C and necessitates UbV^W_dim^ to stabilize one conformation upon binding. (A) Crystal structure of UbV^W^, determined to 2.8 Å resolution, reveals an oligomeric assembly. Two copies of UbV represent the biological unit, colored in orange (UbV^W^ protomer A) and blue (UbV^W^ protomer B). (B) Summary of the results of biophysical analyses of purified UbV mutants and APC2 WHB domain. Wild-type and mutant versions of UbV^W^ were analyzed by Sedimentation Velocity Analytical Ultracentrifugation (AUC) and Size Exclusion Chromatography with Multi-Angle static Light Scattering (SEC-MALS) either as purified apo proteins, or in combination with APC2 WHB domain. (C) Top left, isolated homo-dimer of UbV^W^ from the oligomeric assembly showing protomer A (orange) and protomer B (blue) interacting as a β-strand swapped dimer, representing the biological unit. Bottom left, a composite “monomer” of the UbV^W^ structure superimposes well with Ubiquitin. Notably, the dimer contains two copies of residues shown to be involved in interaction with APC/C, including Trp10 and Ile44 (right). (D) Comparison of UbV^W^ with previously reported domain-swapped dimeric UbVs (UbV.XR, UbV.15.D) reveals a structural relationship; each harbors mutations to amino acid 10 which enables strand-swapping and conformational flexibility between the protomers. Top, UbV from this study with protomer A in orange and protomer B in blue. Middle, published UbV.XR shown in light [PDB: 5O6S] or medium gray [PDB: 5O6T] when aligned over protomer A. Bottom, published UbV.15.D [PDB: 6DJ9] in dark gray when aligned over protomer B. Arrows traces path of alpha helix within protomer B. (E) Selected mutants of UbV^W^ are assessed for association with an immobilized GST-APC2 WHB with purifed GST alone serving as negative control. Asterisk denotes degradation product from residual TEV protease in UbV^W_dim^ prep. (F) Model demonstrating dynamic association between two copies of UbV^W^ per APC/C (left), or stable association of UbV^W_dim^ harboring a single Ile44 mutation per dimer, rendering one copy inefficient at binding APC/C.

Interestingly, the domain swapping of UbV^W^ is reminiscent of that recently reported for other UbVs, one which binds the dimeric RING-RING assembly from the E3 XIAP (UbV.XR) and another which binds the DUSP domain of the DUB USP15 (UbV.15.D) (42, 49). Conformational differences between two independent crystal structures of UbV.XR suggested flexibility in the beta-sheet formed by the domain swap, with different relative orientations varying by 30° for the two halves of the dimer. Superimposition of one monomer of UbV^W^ reveals yet a different arrangement, raising the possibility that domain-swapped UbVs will generally display flexibility across the dimer interface (Fig. 3d).

The NMR spectrum of UbV^W^ was examined for potential dynamic behavior. Indeed, extreme line broadening and multiple peaks indicative of slow-exchange between multiple interconverting conformations precluded resonance assignment for UbV^W^ alone. Nonetheless, assignments of ^13^C/^15^N UbV^W^ in the presence of the APC2 WHB domain and vice-versa enabled identifying intermolecular NOE data, which suggested that the interaction centers around UbV^W^ residues corresponding to Ub’s canonical hydrophobic patch and the UBE2C-binding site on the APC2 WHB. However, residual conformational dynamics precluded structure calculation for the UbV^W^-APC2 WHB complex.

To determine if features of the domain-swap were essential for the interaction, we examined sequences of the known UbVs displaying domain swaps. Notably, UbV^W^, UbV.XR, and UbV.15.D harbor replacements for Ub’s residue Gly10, raising the possibility that Gly10 is important for maintaining a monomeric Ub fold structure. Reverting Trp10 to Gly indeed yielded a monomer in solution as determined by AUC and SEC-MALS (Fig. 3b, Supplemental Fig. 1a, 1b), but also eliminated the interaction with APC2’s WHB domain (Fig. 3e). Although more conservative replacements for Trp10 to Phe, Tyr, His, or Met failed to prevent oligomerization, His and Met substitutions did block assembly with APC2 (Fig. 3b, Supplemental Fig. 1c). Thus, Trp10 is not only essential for the domain swap, but apparently plays a role in mediating UbV^W^ binding to APC2.

To facilitate structure determination, we sought to generate a more uniform domain-swapped UbV that retains Trp10, adopts a dimer without higher-order self-association, and forms a singular assembly with the APC2 WHB domain. An I44D mutant UbV satisfied the criterion of forming a dimer, presumably maintaining the strand-swap and ablating further self-association, but did not bind APC2 (Fig. 3b, 3e). However, following coexpression of a His-tagged UbV^W^ with a GST-tagged I44D mutant, tandem affinity chromatography yielded a heterodimeric domain-swapped dimer that binds APC2 with improved properties (Fig. 3b, 3e, 3f). We term this engineered dimer UbV^W_dim^, with protomer “A” retaining Ile44 and protomer “B” harboring the I44D mutation.

### Structural mechanism of UbV inhibition of APC/C substrate ubiquitylation by UBE2C

UbV^W^-^dim^ retains the original interaction with the APC2 WHB domain, as indicated by nearly identical chemical shift perturbations when UbV^W^ and UbV^W^-^dim^ are added to ^15^N-labeled APC2 WHB domain (Fig. 4a, Supplemental Fig. 2a), and its ability to inhibit UBE2C-mediated ubiquitylation and bind APC2 WHB directly, albeit with reduced affinity attributable in part to the now defunct protomer B (Supplemental Fig. 2b-c). Furthermore, the improved NMR spectral properties enabled structure determination for the UbV^W^-^dim^ complex with APC2 WHB (Fig. 4b, Supplemental Fig. 2e-f, Supplemental Table 2). Representative structures converge to show the same interactions between one Ub-like fold, or “monomer”, of UbV^W^-^dim^ and the APC2 WHB domain, with variable placement of the second Ub-like fold in the domain-swapped dimer (Fig. 4c). Flexibility is apparently imparted by the residues surrounding and including Val8’ to Ile13’ (‘ refers to side-chains from protomer B), as this region displayed particularly broad spectra and the Cα plot for this region differs from that of Ubiquitin (Supplemental Fig. 2g). With the exception of the side chains of Val 8 and Trp 10 located in the strand-swap, this latter Ub-like fold of the dimer does not participate in the interaction and is not discussed further. Rather, we focus on the convergent interactions between one Ub-like fold domain from UbV^W^-^dim^ and APC2, which bury ~700 Å^2^ of total surface area.

**Figure 4.**
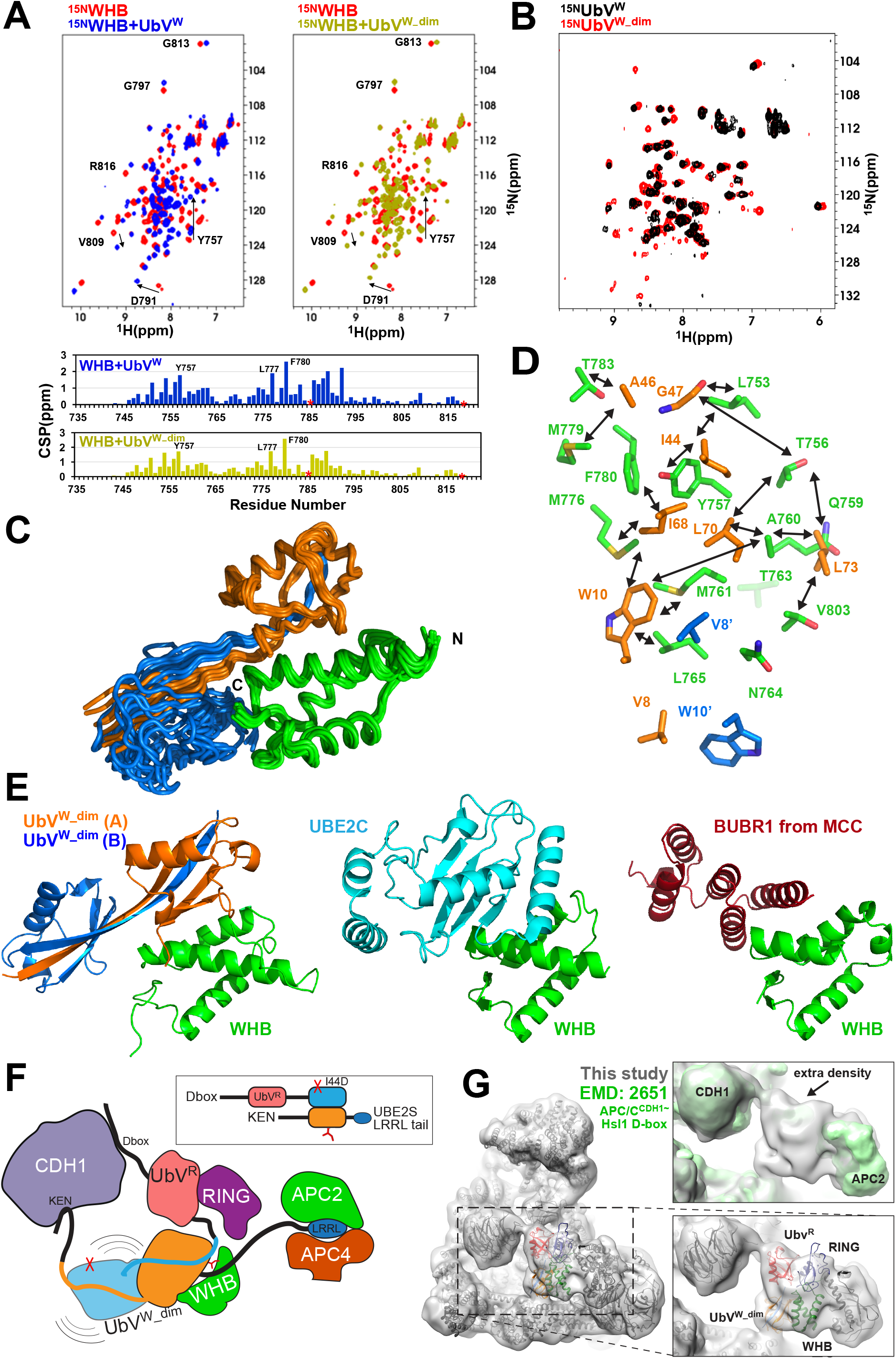
Structure of UbV^W_dim^ bound to APC2 WHB domain reveals molecular basis for blocking UBE2C. (A) Left, overlaid [^15^N, ^1^H] HSQC spectra of ^15^N-labeled APC2 WHB alone (100μM, red) or with UbV^W^ (1:2, blue) with corresponding CSP analysis shown below (blue). Right, overlaid [^15^N, ^1^H] HSQC spectra of ^15^N-labeled APC2 WHB alone (100μM, red) or with UbV^W_dim^ (1:2, gold) with corresponding CSP analysis shown below (gold). Bottom, plot of CSP’s (bar plots) for UbV^W^ and UbV^W_dim^ versus residue number. These results suggest similar interactions between APC2 WHB and the two UbVs. Proline residues indicated with asterisk. (B) Overlaid 2D [^15^N, ^1^H] HSQC spectra of UbV^W^ (black) and UbV^W_dim^ (red) at 308K. UbV^W_dim^ has more resonances resolvable than UbV^W^ illustrating that this sample is more amenable to study by NMR. (C) Solution structures of UbV^W_dim^ bound to APC2 WHB confirm a domain-swapped dimer in solution, composed of ~67% of models that satisfy experimentally determined NOE constraints. All observed conformations of UbV^W_dim^ in the complex differ from all three previously observed crystal structures of domain-swapped dimer UbVs (UbV^W^, UbV.XR, and UbV.15.D), confirming that hinge residues between the Protomer A (orange) and Protomer B (blue) imparts flexibility. However, all Ub-like fold “monomers” superimpose well with Ub (~1.3Å RMSD over Cα) and all populations show the same interaction between Protomer A and the APC2 WHB domain. (D) Residues involved in the interaction between UbV^W_dim^ (orange and blue) and APC2 WHB (green) are illustrated as side chains originating from the ribbon structure. Arrows depict observed intermolecular NOEs in NMR experiments. (E) Model representation of the three observed interactions for APC2 WHB domain. Left, representative structure of UbV^W_dim^ bound to APC2 WHB, compared with prior structures of UBE2C bound to APC2 WHB [PDB: 4YII] (left), and BUBR1 of the Mitotic Checkpoint Complex (MCC) bound to APC2 WHB from the MCC complex with APC/C^CDC20^ [PDB: 5LCW, 5KHU] (right). (F) Graphical illustration of engineered UbV^W_dim^ “trap” designed to associate avidly with many components of APC/C. Coexpression of extended protomers of the UbV^W_dim^ enables multiple points of contact with APC/C. Protomer B of UbV^W_dim^ harbors an N-terminal D-box for recruitment to CDH1 and APC10, and a subsequent UbV^R^ moiety to bind APC11 RING domain. Protomer A of UbV^W_dim^ protomer has an N-terminal KEN-box to bind CDH1, and a C-terminal UBE2S-based extension for docking to APC/C platform subunits. (G) Left, ~9Å lowpass-filtered representation of cryo-EM map of human APC/C^CDH1^ bound to the avid UbV^W_dim^ trap illustrates how UbV^W_dim^ associates with APC2 WHB. Top right, the cryo-EM map is overlaid with a prior map of APC/C^CDH1^-Hsl1 D-box low-pass filtered to similar resolution [EMDB: 2651, green] reveals extra density. Bottom right, docking of structures of UbV^W^ bound to APC2 WHB (orange and green, respectively) and UbV^R^ bound to APC11 RING (red and purple, respectively, PDB: 5JG6) shows how the additional EM density can be attributed to the avid UbV^W_dim^ trap.

As with many complexes with Ub, the interaction is largely hydrophobic, with intermolecular NOEs centering around the UbV^W^ residues corresponding to Ub’s canonical hydrophobic patch (here, Val8, Ile44, Ile68, and Leu70) and the first and second helices and intervening loop of APC2’s WHB domain (Fig. 4d). One end of the complex is secured by the UbV^W^ loop comprising Ala46 and Gly47 inserting in a hydrophobic groove between APC2 helices lined by Leu 753, Tyr757, Phe780, Thr783, and Val803. The other edge is anchored by Leu73, Arg74, and Trp10 enwrapping the C-terminus of the first APC2 WHB helix, through contacts with APC2’s Gln759, Ala760, Met761, Thr763, and Asn764.

Many residues from the UbV^W^-binding surface on APC2 were previously identified as contributing to the recruitment of UBE2C during substrate ubiquitylation, accounting for the inhibition of this reaction (Fig. 2b, 4e). Thus, the structure indicates that UbV^W^ inhibits substrate ubiquitylation by enveloping the surface of APC2 WHB responsible for recruiting UBE2C, competing with the binding of this E2 (27). Interestingly, 55% of APC2 residues contacted by the UbV^W^ also bind to the Mitotic Checkpoint Complex (MCC) in an APC/C^CDC20^-MCC configuration that inhibits ubiquitylation during the spindle assembly checkpoint, prior to proper chromosome alignment on the mitotic spindle (47, 48) (Fig. 4d, 4e).

We wished to confirm the UbV^W^-^dim^ binding site within the full APC/C complex. However, previous studies showed that mobility of the APC2 WHB domain precludes its visualization in cryo EM maps of APC/C, unless harnessed through multisite interactions (reviewed in (50)). For example, the APC2 WHB domain was best visualized in APC/C^CDC20^ bound to MCC, which naturally restricts flexibility through numerous interactions with multiple subunits in APC/C^CDC20^ (47, 48). It was also possible to visualize APC2’s WHB domain in a structure representing substrate ubiquitylation, where an avid multisite binder mimicking the UBE2C-Ub-substrate intermediate was generated by chemistry and protein engineering (27). On this basis, we engineered a complex in which UbV^W^-^dim^ was appended to additional sequences to avidly capture multiple mobile sites on APC/C^CDH1^ and visualize the interactions with APC2. Briefly, a KEN-box sequence from the substrate Hsl1 was encoded N-terminal of the original UbV sequence in UbV^W^-^dim^, while a C-terminal UBE2S recruitment sequence would further anchor this protomer via interactions with the APC/C platform. The I44D mutant protomer B of UbV^W^-^dim^ was fused at the C-terminus of a construct harboring the high-affinity D-box sequence from Hsl1 followed by the RING-binding UbV^R^. (Fig. 4f). This engineered UbV^W^-^dim^ complex, also containing multiple substrate and APC/C binding elements, potently inhibits both UBE2C- and UBE2S-mediated APC/C reactions (Supplemental Fig. 2d) and by using it to purify recombinant APC/C^CDH1^, we obtained a cryo EM density map at an overall resolution of 6.6 Å. The overall map clearly resolves the majority of secondary structures, enabling fitting the corresponding regions of prior high-resolution structures of APC/C and CDH1. As expected, the APC2 WHB and APC11 RING domains were less well resolved. However, extra density corresponding to these domains and their associated UbVs is clearly observed upon downsampling the EM map to 9 Å and comparing with the prior map of an APC/C^CDH1^ complex with the Hsl1 D-box peptide (25) in which the APC2 WHB and APC11 RING domains were not readily visible at this resolution (Fig. 4g). Although the resolution, together with similarity in the sizes of the two UbV complexes, precludes their definitive docking, it is possible to approximately place the structures of APC11 RING-UbV^R^ and APC2 WHB-UbV^W^ based on physical limitations imposed by their covalent linkage to other domains that are clearly resolved in the map. On this basis, the EM data show how the UbV^W^ can block UBE2C binding in the context of the fully-assembled APC/C^CDH1^ complex (Fig. 4g). Notably, our engineered inhibitor acts much like the natural inhibitor MCC, by hijacking and reorienting APC2’s mobile WHB domain.

### APC2 WHB is a ubiquitin-binding domain

Because our prior studies of E3-targeting UbVs selected for tighter binders to known Ub-interacting sites on RING and HECT domains (21, 28, 42), we considered the converse possibility: could UbV binding identify an unknown binding site on an E3? This question was addressed by NMR assays, whereby increasing concentrations of unlabeled Ub was titrated into a ^15^N-labeled version of the isolated APC2 WHB domain, and vice-versa. Indeed, ^15^N-^1^H HSQC spectra showed that addition of high concentrations of Ub (500μM titrant shown) caused chemical shift perturbations at the UbV-binding surface of APC2’s WHB domain (Fig. 5a). Concordantly, the APC2 WHB domain elicited chemical shift perturbations of the hydrophobic surface on Ub (Supplemental Fig. 3a,b). Thus, the APC2 WHB domain is a Ub-binding domain.

**Figure 5.**
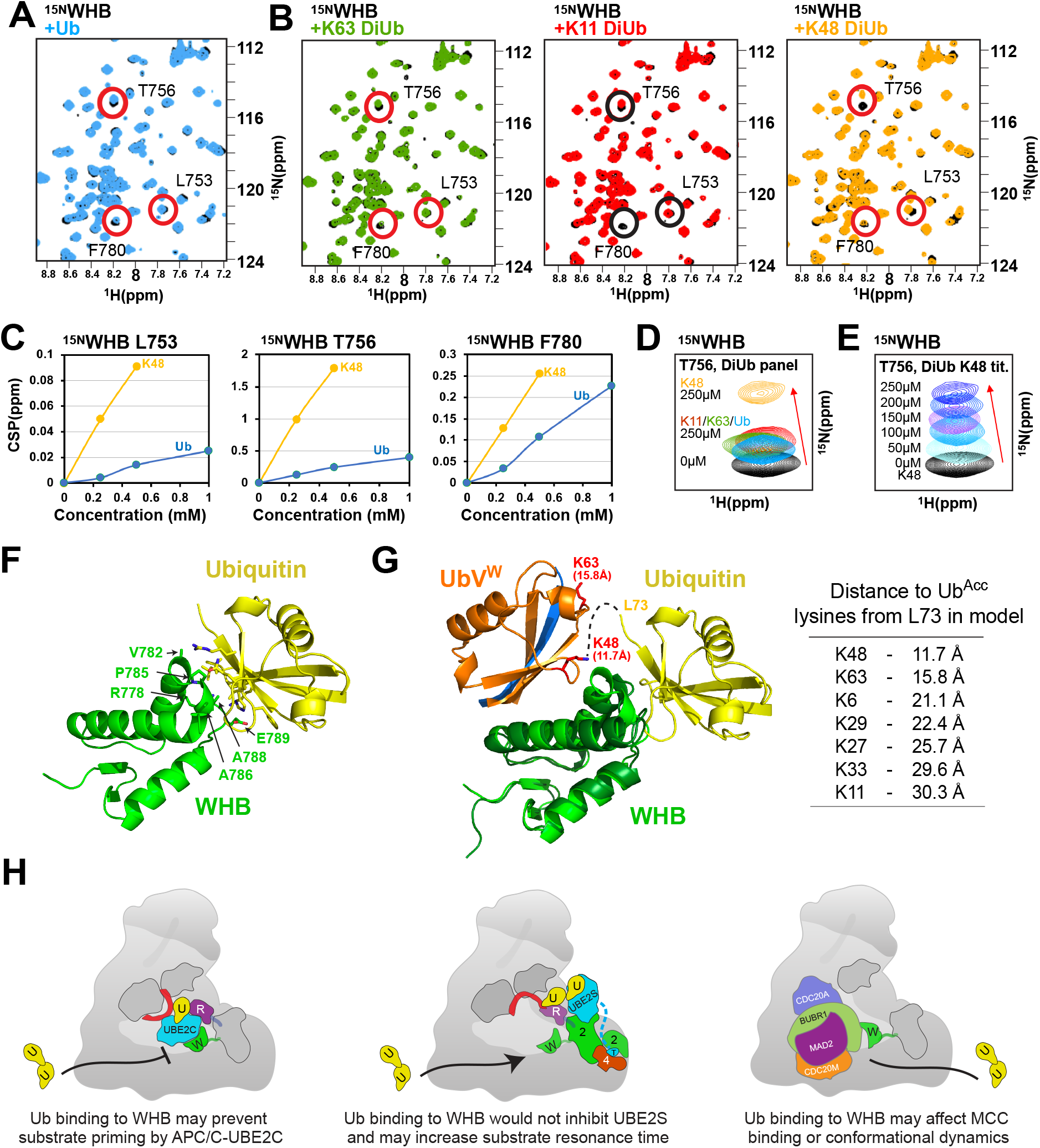
APC2 WHB domain binds Ubiquitin conjugates linked by K48. (A) Overlaid [^15^N, ^1^H] HSQC spectra of ^15^N-labeled APC2 WHB alone (200μM, black) or with Ubiquitin (1:2.5, light blue). Ubiquitin itself causes minor chemical shifts of the APC2 WHB spectra, including for residues L753, T756, and F780 which associate with UbV^W^ in Fig. 4. (B) Overlaid [^15^N, ^1^H] HSQC spectra of ^15^N-labeled APC2 WHB alone (200μM, black) or with purified, enzymatically assembled diubiquitins of distinct linkages. Left, K63 diubiquitin, 250μM, green. Middle, K11 diubiquitin, 250μM, red. Right, K48 diubiquitin, 250μM, gold. (C) CSP values at the indicated residues are plotted for titrations of K48-linked diubiquitin and purified Ubiquitin. Titration of K48 diubiquitin causes a greater chemical shift than Ubiquitin alone. (D) Chemical shift perturbations observed for residue T756 at the Ubiquitin-APC/C interface for different forms of Ubiquitin. The titrants and concentrations are marked. (E) Chemical shift perturbations observed for residue T756 at the Ubiquitin-APC/C interface for titrations of K48-linked diubiquitin. The titrant concentrations are marked. (F) 2.2Å Crystal structure of Ubiquitin bound to isolated APC2 WHB domain. Distinct residues of APC2 are observed associating with Ubiquitin, including V782, P785, R778, E789, A788, and A786. (G) Left, superimposition over the APC2 WHB domain of the crystal structure and solution structures reveals two distinct locations for Ub-like molecules to associate with APC. Dashed lines indicate a possible trajectory for 3 remaining C-terminal residues of Ubiquitin to form a K48 linked chain. Right, distances to each side-chain Lysine of Ubiquitin are measured from Leu73 of the donor Ubiquitin in the shown model and listed ascendingly. (H) Cartoon model demonstrating potential functional consequences of Ub~WHB interaction. Left, Ub chain association with WHB could compete for UBE2C recruitment to APC/C, inhibiting ubiquitylation of substrate lysines. Middle, Ub recruitment to APC2 WHB could enhance UBE2S activities in multiple ways: here, substrates are retained longer on APC/C, while UBE2S would also have unhindered access to APC/C with the UBE2C-binding site blocked. Right, Ub association with WHB could have mutliple potential effects on MCC association with APC/C. Ub chain may compete with BUBR1 for contacting WHB. Disrupting the MCC-WHB interaction could lead to an alternate conformation or aid in MCC dissociation from APC/C. Alternatively, harboring Ub that is conjugated directly to MCC could further affect the conformational landscape and limit dissociation of the inhibitor itself.

### APC2 WHB preferentially binds K48-linked ubiquitin chains

Does APC2’s WHB domain preferentially bind a particular type of Ub chain? To address this question, we first purified linkage-specific di-Ub conjugates (Supplemental Fig. 3c) for three architectures: K11 and K48-chains, based on human APC/C producing these linkages, and K63-chains, which are not recognizable components of APC/C-dependent signaling but which are highly abundant in cells. Chemical shift perturbations were monitored in ^15^N-^1^H HSQC spectra of the ^15^N-labeled APC2 WHB domain upon titrating each of these chains (Fig. 5b), but only the conjugates harboring a K48-linkage elicited greater chemical shift perturbations relative to the effects of Ub alone (Fig. 5c-e, Supplemental Fig. 3d). Notably, the isolated RING domain exhibited no preference for any chain linkage type under identical experimental conditions (Supplemental Fig. 3e-g) highlighting the specificity of K48 diubiquitin’s association with the APC2 WHB domain.

We wished to obtain insights into how K48-linked di-Ub might preferentially bind the APC2 WHB domain, although we were unable to obtain a structure of a complex with a Ub chain, possibly due to intramolecular competition for WHB binding or a slight collateral restructuring of Ub-binding sites evidenced in Cα deviations for this region upon UbV^W^ titration (Supplemental Fig. 4a). It seems likely that one Ub binds in a manner similar to the UbV^W^-^dim^, because the few resonances that shift upon binding the K48-linked di-Ub all correspond to residues at the interface in the APC2-UbV^W^-^dim^ complex. Thus, a potential placement for one Ub can be modeled by grafting Ub in place of the main Ub-fold from UbV^W^-^dim^. A possible model for the second Ub was provided by a fortuitous 2.2 Å resolution crystal structure of a single Ub bound to APC2’s WHB domain (Fig. 5f, Supplemental Table 1). In the crystal, Ub’s C-terminus points toward the UbV/E2 binding surface, and its hydrophobic patch mediates interactions via an APC2 surface involving Arg778, Val782, Pro785, Ala786, and Ala788, and Glu789. This interaction differs from the main contacts revealed by NMR and is presumably stabilized by crystal contacts supported by the high protein concentrations that occur during crystallization. Notably, the primary Ub binding site observed by NMR is shielded by crystal contacts. Nonetheless, the distances measured between the C-terminal residue visible in the crystal (Leu73) and each lysine in the modeled primary-binding Ub provide a potential rationale for the linkage preference for conjugate binding: only a K48-linked Ub chain would be compatible with this di-Ub arrangement (Fig. 5g). Consistent with our hypothesis, ubiquitylation of substrate (Supplemental Fig. 4b) leads to a less productive E2 encounter monitored by the rate of discharge of Ub from the catalytic cysteine of UBE2C in the presence of APC/C^CDH1^ (Supplemental Fig. 4c), although future studies will be required to definitively determine the basis for this reduced activity, the structural details for K48-linked Ub chain binding to APC2, and the functions of these interactions.

## DISCUSSION

UbVs are emerging as powerful tools for probing functions of the Ub system. To date, we and others have selected for UbVs that bind catalytic domains from enzymes (DUBs, E3 ligases, E2 conjugating enzymes), interchangeable cullin-binding subcomplexes (Skp1-F-box proteins), and Ub-binding domains (21, 28, 38–44). These selections have yielded hundreds of tools targeting the known domains by modulating catalytic activity and assembly of E3 ligases, and inhibiting Ub binding by well-recognized interacting domains. Here, we identified a UbV modulating activity of the massive, multifunctional RING E3 ligase, APC/C, through an entirely different route: scanning through a panel of purified UbVs for effects on in vitro ubiquitylation (Fig. 1b). Remarkably, even a relatively small collection of UbVs contained one (UbV^W^) with a novel mode-of-action, selectively inhibiting ubiquitylation by only one of the two E2s associated with APC/C. Importantly, UbV^W^ also inhibits APC/C-dependent substrate turnover in the ex vivo setting of mitotic extracts from *Xenopus* eggs (Fig. 1f). Mechanistic and structural studies revealed that unlike other characterized UbV inhibitors of ubiquitylation, UbV^W^ does not bind a hallmark E3 ligase catalytic domain. Rather, UbV^W^ blocks an auxiliary site - the WHB domain from APC/C’s APC2 subunit - from selecting and positioning UBE2C for substrate priming (Fig. 2, 4). E2 specificity was achieved because this APC2 domain is dispensible for APC/C’s other partner E2, UBE2S provided that substrates are pre-marked with a Ub for chain elongation. Interestingly, an emerging theme in E2-E3 interactions is the importance of secondary interactions beyond hallmark RING domains in establishing specificity. Thus, our results raise the possibility that UbVs may generally be useful tools for targeting such auxiliary interactions to selectively modulate only subsets of E3 activities. In terms of UbV^W^, we envision that more directed studies in the future could be used to select for tighter-binding UbVs that target APC’s WHB domain, which could serve as affinity probes for cell-based studies aimed at understanding biological roles of UBE2C.

UbV^W^ also guided our unanticipated identification of APC2’s WHB domain as a novel Ub-binding domain in APC/C (Fig. 5). Reasoning that UbVs preserve many key features of the Ub fold, and that the majority of UbVs studied to date interact with known Ub-binding sites, we tested for Ub-binding by NMR. The limited chemical shift changes at high protein concentrations likely reflect extremely weak interactions. Such weak binding is on the same scale as Ub interactions with other well-characterized, validated partners. This includes APC11’s RING domain, which recruits a free Ub with a *K_m_* in the millimolar range for UBE2S-dependent K11-linked chain formation (22). It is thought that weak Ub interactions are important for contributing as one of several elements to multisite interactions (36, 37). Several protein-protein interactions that are weak on their own can synergize to produce high local concentrations sufficient for binding with high specificity. Such interactions are also dynamic, because binding is achieved through avidity of many simultaneous interactions, while disruption of any individual interaction could be sufficient to dismantle an assembly. Another explanation for low affinity is to prevent errantly blocking an individual Ub-binding site by the massive amount of Ub present in non-cognate contexts.

In terms of Ub binding to APC2’s WHB domain, several scenarios could establish avidity. In addition to K48-linked chains that could bind to APC2, ubiquitylated APC/C substrates harbor one or more distinct degrons interacting with a coactivator (reviewed in (51)). Ubiquitylation could also modulate binding of APC/C regulators, such as Mitotic Checkpoint Complex. Another source of avidity may arise from multiple Ubs linked to each other, and/or to different sites on substrates. Indeed, a K48-linked conjugate (i.e. harboring two Ubs) shows increased interaction with APC2’s WHB domain (Fig. 5c-d), while additional substrate-linked Ubs could potentially capture the Ub-binding exosite on APC11’s RING domain (22, 28). Notably, our cryo EM map enabling showing APC/C interactions with UbV^W_dim^ relied on conceptually related multisite interactions also involving two substrate degrons and two APC/C-binding elements (Fig. 4g).

As with many functionally important Ub-binding exosites upon their initial discovery (52–54), the role(s) of Ub binding to APC2’s WHB domain remain unknown. Although the requirement for this same surface for substrate ubiquitylation imposes technical challenges for studying function, it also implies potential roles, either antagonizing UBE2C, or participating in UBE2C-independent activities. One such activity is polyubiquitylation by UBE2S after UBE2C-dependent priming. In principle, APC2’s WHB domain could capture emerging K48-linked chains on substrates, increasing a substrate’s lifetime on APC/C and thereby potential for UBE2S-dependent modification with K11-linked chains. This could also contribute to switching between the two E2 activities, or modulating binding of Mitotic Checkpoint Complex (Fig. 5h). Although it is presently unknown whether UBE2C and UBE2S can function simultaneously or not on a single APC/C complex, if ubiquitylation by the two E2s is mutually exclusive, then Ub binding to APC2’s WHB domain would block UBE2C and thereby enable K11-linked chain formation by UBE2S. APC2’s WHB domain could also contribute to localizing APC/C to particular subcellular localizations harboring K48-linked chains or chain-forming activity. It is compelling that these mechanisms could all contribute to marking of substrates with both K48 and K11 linkages, a feature of APC/C substrates recently shown to direct their 26S Proteasomal degradation (33, 55).

Finally, our study demonstrates that UbV technology can identify novel Ub-binding sites within massive multiprotein complexes. As UbVs are traditionally selected in binding assays, much like the generation of other affinity reagents through phage- or other display technologies, it seems likely that use of entire assemblies even in the megadalton range could be used to discover UbVs. However, UbV^W^ was instead identified through an enzymatic assay (Fig. 1b). Thus, extending the modalities by which UbVs could be expressed and selected, for example through expression in pools, could be useful for identifying UbVs that mimic myriad functions of Ub binding. We envision numerous ways one could affect in vitro ubiquitylation depending on the order of addition of reaction components, including through modulating multiprotein complex E3 assembly, substrate binding, E2 binding, Ub ligation, ubiquitylated substrate binding, and linkage specificity of polyubiquitylation. Such approaches may be particularly useful for identifying cryptic Ub-binding sites regulating dynamic multifunctional E3s like APC/C. Although we have now identified several exosites – on the APC11 RING domain, and on the APC2 WHB domain – these do not completely eliminate processivity of multi and polyubiquitylation by UBE2C, indicating that additional Ub-binding sites on APC/C remain to be discovered (22, 28, 34, 35). It seems like other massive molecular machines in the Ub system, including the 26S Proteasome, many E3s and their associated E2s will emerge to be highly regulated through numerous Ub-binding sites acting in concert and yet dynamically to modulate function (5, 7, 37). We thus anticipate massive capabilities for UbVs to both unearth and target these sites to transform our understanding of roles of Ub in regulation.

## MATERIALS AND METHODS

Recombinant APC/C, CDH1, and UBA1 were expressed in High-five insect cells, and all other proteins in Escherichia coli as decribed previously (22,28). The complex “trap” for visualizing UbV^W^ binding to APC2 WHB in the context of the full APC/C^CDH1^ complex was the product of coexpression of two gene products encoding several APC/C^CDH1^ binding moeities; more details are available in the SI. Ubiquitylation reactions, NMR, and substrate degradation assays in X. laevis egg extracts were largely performed as described previously 22,28). Crystallographic data were collected at Advanced Photon Source (APS) Northeastern Collaborative Access Team (NECAT) ID-24-C and Southeast Regional Collaborative Access Team (SERCAT) ID-22 beamlines. Further details can be found in the SI.

## Supporting information

Full supplement as single pdf

## Author contributions

E.R.W., C.R.R.G., D.J.M, N.G.B., H.S., J.-M.P., S.S.S., and B.A.S. designed research; E.R.W, C.R.R.G., D.J.M., W.Z., I.F.D., J.R.P., D.H., S.Y., D.L.B., E.T.K., R.V., and N.G.B. performed research; E.R.W., C.R.R.G., W.Z., D.J.M, I.F.D., J.R.P., N.G.B., H.S., J.-M.P., S.S.S., and B.A.S. contributed new reagents/analytic tools; E.R.W., C.R.R.G., D.J.M, J.R.P., N.G.B., J.-M.P., S.S.S., and B.A.S. analyzed data; and E.R.W., N.G.B., C.R.R.G., D.J.M., and B.A.S. wrote the paper.

## Acknowledgements

We acknowledge Dr. J. Kellermann, Dr. S. Uebel, Dr. K-P. Wu, M. Zobawa, and M. Brunner for their scientific and technical advice. We are grateful to Dr. M. Strauss for training and assistance with the cryo EM experiments. We thank Dr. V. Chau and Dr. C. Coleman for the GP78-UBC7 plasmid used for the generation of K48-linked diubiquitin. For funding we thank CRS scholarship for the next generation of scientists (W.Z.); Deutsche Forschungsgemeinschaft Sonderforschungsbereich 860 (H.S.); Boehringer Ingelheim, the Austrian Research Promotion Agency (FFG Laura Bassi Centre for Optimized Structural Studies), the European Union (Seventh Framework Programme Grant 227764 MitoSys), and the Austrian Science Fund (SFB-F34 and Wittgenstein award) (J.-M.P.); NIH R35GM128855 and UCRF (to N.G.B.); Genome Canada DIG grant OGI-119 (S.S.S.); and ALSAC, HHMI, NIH R37GM065930 and P30CA021765 (BAS).

