## Supplementary material for "A Protein Engineering Approach for Uncovering Cryptic Ubiquitin-binding Sites: from a Ubiquitin-Variant Inhibitor of APC/C to K48 Chain Binding": Full supplement as single pdf

### Supplemental Experimental Procedures

**Purification of proteins for enzyme assays, binding studies, NMR, and crystallography.** For ubiquitylation assays, human APC/C, CDH1, UBA1, UBE2C, UBE2S, Cyclin B (residues 1-95), and UbCyclin B (residues 1-95) were purified as previously described (1, 2). Recombinant APC/C was expressed in a baculovirus expression system with a C-terminal twin-Strep tag on APC4, similar to recent structural studies (3, 4). For all experiments other than EM, APC/C was purified with a 3-step scheme: Affinity purification with Strep-Tactin Sepharose (IBA Lifesciences) and elution with desthiobiotin, anion exchange with gradient NaCl elution, and size exclusion chromatography (SEC) in a final buffer of 20 mM HEPES pH 8, 200 mM NaCl, 1 mM DTT. APC2 WHB domain (residues 735-822) was expressed in BL21(DE3) Gold cells from a modified pGEX 4T1 vector encoding a TEV-cleavable GST tag at the N-terminus. Following GST-affinity and on-column cleavage, WHB in the flow-through was treated to anion exchange with gradient NaCl elution, and size exclusion chromatography (SEC) in a final buffer of 20 mM HEPES pH 8, 200 mM NaCl, 1 mM DTT.

N-terminal 3x MYC-$His_6$-HRV14 3C protease site-CDH1 was expressed in a baculovirus expression system and purified by nickel-affinity chromatography. After HRV14 3C-mediated proteolytic cleavage of the 3X MYC-$His_6$ tag, CDH1 was then subjected to cation exchange chromatography and SEC in a final buffer of 20 mM HEPES pH 7, 300 mM Ammonium Sulfate, 1 mM DTT, 2.5% Glycerol.

UBE2C, UBE2S, and gp78-Ubc7 fusion were expressed in BL21(DE3) Codon Plus (RIL) cells. UBE2C was isolated by nickel-affinity chromatography via a C-terminal $His_6$ tag and purified by SEC in 20 mM HEPES pH 8, 200 mM NaCl, 1 mM DTT. UBE2S was expressed in a modified pRSF duet vector with a $His_6$-TEV protease site-FLAG-HRV14 3C protease site-fused to the N-terminus, isolated by nickel-affinity chromatography, subsequently cleaved by HRV14 3C protease, further purified by cation exchange chromatography, and finally purified by SEC in 20 mM HEPES pH 8, 200 mM NaCl, 1 mM DTT. Hybrid gp78-Ubc7 E2 fusion was isolated by nickel-affinity chromatography on an N-terminal $His_6$ tag and further purified by anion exchange chromatography and SEC in 20mM Tris pH 7.5, 150mM NaCl, 1mM DTT.

Cyclin B 1-95 (CycBNT*) and Ub-Cyclin B 1-95 (UbCycBNT*) were purified as previously described (1, 5). In short, these substrates were expressed with an N-terminal GST-TEV and C-terminal Cys-$His_6$ fusion in BL21(DE3) Codon Plus (RIL) cells. The substrates were then purified by glutathione-affinity chromatography, TEV-mediated proteolysis, and nickel-affinity chromatography. The substrates were then fluorescently labeled (denoted by an asterisk) as previously described (2), with fluorescein-5-maleimide. Subsequent buffer exchange via Zeba spin columns and SEC was used to remove the excess fluorescein.

Ubiquitin variants (UbV) used as inhibitors were expressed in BL21(DE3) Codon Plus (RIL) cells, isolated by an N-terminal $His_6$-tag, and normalized for quality by coommassie-brilliant-blue stained SDS-PAGE. $UbV^{W}$ and the $UbV^{neg}$ mutant used in assays monitoring degradation of APC/C substrates in Xenopus Egg Extracts were subjected to TEV-mediated proteolytic cleavage and SEC in a final buffer of 20 mM HEPES pH 8, 200 mM NaCl. $UbV^{W\_dim}$ was generated by co-transformation of a pRSF $His_6$-TEV UbV WT and a pGEX GST-TEV UbV (I44D) for co-expression in BL21(DE3) Codon Plus (RIL) cells. Protein was isolated first by GST-affinity chromatography and selected for homogeneity by subsequent nickel-affinity chromatography. Following TEV-mediated cleavage of both tags and anion exchange chromatography/GST-affinity clearing the GST-tag and uncleaved contaminants, final stoichiometric purity was achieved by SEC and validated by intact mass spectrometry.

Diubiquitin chains were generated enzymatically according to published protocols with minor differences (6, 7). In brief, for the generation of K11-linkages, a K11-specific Ube2s-Uba fusion was used as E2 enzyme (similar to addgene plasmid #66713; http://n2t.net/addgene:66713) and reactions were supplemented with AMSH to counteract non-specific K63 linkages. For the generation of K48-linkages, a K48-specific gp78-Ubc7 fusion was used as E2 enzyme and both AMSH and Cezanne were used to counteract non-specific K63 or K11 linkages. For the generation of K63 linkages, The Ubc13-Uev1a heterodimer was used as E2 enzyme. Reactions consisted of 1-10 μM E2 enzyme, 1μM Uba1, 0.5 μM deubiquitylating enzyme, 10 mM ATP/MgCl2, 40 mM Tris pH 7.6, 0.6 mM DTT, and 6 mM Ubiquitin. Excess Ubiquitin is used to direct the production of diubiquitin and small chains. Following overnight incubation at 37°, reactions were quenched with the addition of 50 mM DTT and diluted 15-fold with 50 mM Ammonium Acetate pH 4.5. Reactions were purified by cation exchange and iterative size exclusion chromatography in a buffer containing 10 mM HEPES pH 7, 100 mM NaCl, 10 mM DTT until homogeneity.

The affinity chomatography assay in Fig. 3e is the product of coexpression of GST-tagged APC2 WHB with $His_6$-tagged versions of the indictated UbV except in the case of $UbV^{W\_dim}$. Since $UbV^{W\_dim}$ is already the product of coexpression, it is added post-cleavage and purification to immobilized GST-WHB, allowed to associate 30 minutes followed by washing and elution in the same format as the neighboring lanes. Gel-loading is normalized for total protein mass. Due to small amounts of contaminating TEV protease from the $UbV^{W\_dim}$ purification, that sample exhibits minimal proteolysis following pulldown/elution and prior to quenching.

Polyubiquitylated Securin ($Ub_n$-Securin) used in Supplemental Fig. 4c, was formed by mixing 25 μM Ub-Securin, 1 μM UBA1, 10 μM UBE2C, 1 μM CDH1, 100 nM APC/C, 10 mM $MgCl_2$-ATP, and 125 μM Ubiquitin for 4 hours at room temperature. $Ub_n$~Securin was purified by nickel affinity chromatography, buffer exchanged into 20 mM HEPES pH 8, 200 mM NaCl, and flash-frozen. Degree of ubiquitylation is visualized via SDS-PAGE and western blot for a C-terminal His tag (Supplemental Fig. 4b).

**Enzyme assays.** Qualitative assays probing the function of APC/C E2s in the presence or absence of UbV proteins were performed with 30 nM APC/C, 500 nM CDH1, 0.2 μM UBE2C or UBE2S (unless otherwise indicated), 10 mM ATP/MgCl2, 0.1 μM E1, 125 μM Ub, 0.2 μM CycBNT* or UbCycBNT* and the indicated concentrations of UbV, except Fig. 1e and 2e, which use 1μM CDH1 and 65μM Ubiquitin, and Fig. 2e receives isolated WHB domain at the indicated concentration. Assays are quenched in 1x-SDS loading dye at the 12 minute timepoint, and fluorescent substrate is visualized using a Typhoon FLA9500 PhosphorImager following SDS-PAGE.

In kinetic experiments, apparent *K*m ($Km^{app}$) and apparent *V*max ($Vmax^{app}$) values were determined by fitting the initial velocities to the hyperbolic Michaelis-Menten, $v = Vmax^{app}[X]/(Km^{app} + [X])$, equation, where X is the UBE2C concentration, using GraphPad Prism 6 software. Single time points were taken under conditions that satisfy initial velocity regimes. In summary, a time course was monitored at both the minimum and maximum point of each titration to ensure a single timepoint could be taken where the substrate depletion is minimal and product formation remained linear.

The kinetic parameters, *K*m$^{app}$ and *V*max$^{app}$ values, were determined for UBE2C (Fig. 2a) in assays monitoring substrate polyubiquitylation with APC/C$^{CDH1}$ to evaluate E2–APC/C binding and the maximum rate of ubiquitylation. Concentrations of 10 nM APC/C, 50 nM E1, 10 mM ATP/MgCl2, 65 μM Ub, 100 μM UbV$^{W}$, and 200 nM UbCycBNT* (residues 1-95) were used and the reactions were quenched after 6 minutes at room-temperature.

Pulse-chase assays, used to examine the ability of the APC/C to activate UBE2C~*Ub were performed by loading fluorescein-labeled ubiquitin (*Ub) to 25 μM UBE2C by the addition of 1 μM UBA1 and 5 mM $MgCl_2$–ATP for 2 minutes, and quenching with 0.5 mM EDTA pH 8. UBE2C~*Ub was then added to 100 nM APC/C, 150 nM CDH1, and either 25 μM $Ub_n$-Securin or unmodified Securin. After SDS-PAGE, a time course of UBE2C~*Ub bands were imaged and quantitated based on the fluorescent Ub using a Typhoon FLA 9500.

**UbV$^{W}$ identification.** The preparation of the UbV phage-displayed library, protein immobilization and following phage selections were done according to established protocols (8). UbV$^{W}$ was not isolated based on their affinity for UBE2C or APC2, rather it was selected from a pool of existing UbVs available in the Sidhu laboratory after testing for inhibitory effects in APC/C-specific assays.

**NMR spectroscopy.** All NMR experiments were performed in 10 mM HEPES pH 7, 100 mM NaCl, 10 mM DTT, 10% $D_2O$ on either a Bruker Avance 600-MHz or 700-MHz spectrometer equipped with a 5-mm triple resonance cryoprobe and a single-axis pulse field gradient. Apo WHB assignments were done by our group earlier (BMRB ID: 26527) and the assignments were further confirmed at 308K using 500 μM $^{15}$N, $^{13}$C-WHB in the same buffer using 3D HNCA. The complexes of $^{15}$N, $^{13}$C-WHB with unlabeled UbV$^{W}$ and UbV$^{W\text{–}dim}$ were reassigned at 298K, 308K and 318K using 3D HNCA. Side chain assignments of the complex were finalized using $^{15}$N-edited NOESY, HCCH-$^{13}$C-edited TOCSY, HCC(CO)HN and $^{13}$C-edited aliphatic and aromatic NOESY spectra (9). The sample behaved much better at higher temperature but was not stable over the long periods of NMR measurement time. Hence, we collected the NOESY spectra at either 308K or 298K for structure calculations.

The 2D [$^{15}$N, $^{1}$H] spectra of apo UbV$^{W}$ and UbV$^{W\text{–}dim}$ showed severe line broadening (Fig. 4b) for several resonances in the β-sheet regions, such that the assignment could not be completed. The backbone resonances were thus assigned only for the complex of $^{15}$N, $^{13}$C- labeled UbV$^{W}$ and UbV$^{W\text{–}dim}$ and unlabeled WHB in a molar ratio of 1:2, using 300 μM sample (Supplemental Fig. 2a, middle, right). This complex exhibited two sets of resonances for most of the peaks in the β-strand region and one broad resonance for the residues in the helix of UbV, as expected from the crystal structure of the domain-swapped dimer. First, we collected the 3D HNCA and HNCO data at 328K using 40% non-uniform sampling. Since the sample was not stable at high temperature, we re-measured the data at 298K and 308K and all the NOESY data were collected also at these two temperatures (Supplemental Fig. 2a). Even in the presence of WHB, several UbV$^{W}$ resonances, mostly in the hinge region showed very broad resonances which include V8/V8', Q9/Q9', W10/W10', K11/K11', T12/T12' and I13/I13'. Even the C$\underline{\alpha}$ resonances (only a very weak peak was present) were broad in the HNCA spectrum, suggesting that the backbone of this region is moving in msec time scale that results in significant line broadening for these resonances. Increasing the sample to high temperatures, 318K (45° C), did not sufficiently increase the signal. While the methyl resonances of V8/V8' and side chain resonances of Q9/Q9' and K11/K11' could be assigned, the backbone resonances were broad and only sequential backbone NOE's were strong and hence were used in the structure calculation around this region. Also, the chemical shifts of the Cα, Cβ resonances were confirmed with the help of C-

HSQC/NOESY peaks observed from the aliphatic carbon NOESY spectra. Based on the Cα shifts that suggest a domain-swapped conformation in this region (Supplemental. Fig. 2g, black) as well the observed αH-NH cross peaks, hydrogen bonds were proposed in this region for the backbone resonances and the crystal structure was used to aid this along with 15N-NOESY spectra (absence of exchange peaks with water resonance). Further bundling of backbone in this region resulted from only intermolecular NOEs with WHB. Thes broadening of resonances around this region is in support of conformational dynamics around this hinge region in the backbone of UbV. Side-chain resonances were assigned from the combined information content of $^{15}$N-edited TOCSY-HSQC, $^{15}$N-edited NOESY, HCCH-$^{13}$C-edited TOCSY, HCC(CO)HN, HBHA(CO)HN spectra (10). NMR data were processed by the Topspin 3.5 software and analyzed by CARA (11). Resonance assignment of $UbV^{W\_dim}$ is deposited in the BioMagResBank.

Except for the Hα protons in the domain swapped region containing residue, D6, T12, A66, and residues around the mutant I44D, L43, F45, most of the side chain resonances of all the residues showed a single peak. Because of this resonance overlap, and using the symmetry from crystal structure, we assigned the resonances of one of the strongest monomer and the observed NOE's for this monomer were used to create the second symmetric monomer. Wherever the second resonance was assigned, they were used to confirm the presence of NOEs across the β-strand in the domain swapped region, guided by the crystal structure. Also, NOEs observed to and from D44' were calibrated and replaced by the I44 NOEs. NOE distance restraints were obtained from $^{15}$N-edited NOESY-HSQC and $^{13}$C-edited aliphatic and aromatic NOESY-HSQC spectra at a mixing time of 120ms. Backbone dihedral angle restraints were predicted by the TALOS+ software (12). Alpha-helical hydrogen bonds were introduced for the amides that were missing water-exchange cross-peaks in $^{15}$N-edited NOESY spectra. Intermolecular NOE's were measured using two different samples (Figure 4d), using $^{13}$C, $^{15}$N-filtered ($\omega_1$) and $^{13}$C-edited NOESY spectra using a mixing time of 120ms. The NOE's from WHB to $UbV^{I44D}$ resonances were measured using $^{13}$C, $^{15}$N-labeled WHB and unlabeled $UbV^{W\_dim}$ complex (Supplemental Fig 2e). The reverse NOE's from $UbV^{W\_dim}$ resonances to WHB were measured using $^{13}$C, $^{15}$N-labeled $UbV^{W\_dim}$ and unlabeled WHB complex (Supplemental Fig 2f). Though the backbone resonance of W10 was broadened due to slow ms-μs motion, the aromatic protons could be assigned without ambiguity, using carbon chemical shifts, even though they exhibited severe line broadening. The aromatic protons of W10 showed several inter-molecular NOE's to the methyl resonances of residues A760, M761 and L765 (Supplemental Fig. 2e). Similarly, the aromatic resonances of Y757 and F780, that showed the largest chemical shift perturbation in WHB, showed intermolecular NOEs to the methyl resonances of I44 and I68 residues of UbV (Supplemental Fig. 2f). The entire network of intermolecular NOEs observed are shown schematically in Fig. 4d. Most of these NOEs were from the methyl resonances of these residues, except G47 of $UbV^{W\_dim}$, which showed backbone NH and H$\underline{\alpha}$ NOE's to the methyl resonances of L753 (Supplemental Fig. 2e). A total number of 57 inter-molecular NOE's were observed between WHB and $UbV^{W\_dim}$ and the statistics of the intramolecular NOE's used in the structure calculation is given in Table S2. Structures were initially calculated by the program UNIO (13), using CYANA for energy minimization, and the final calculations were made manually using CYANA (14). Initially we calculated 400 structures and the lowest energy structure is used as a representative model for structural comparison. Though there is conformational flexibility between $UbV^{D44}$, with respect to the WHB-$UbV^{I44}$ complex, mainly due to absence of NOEs, this monomer by itself if overlapped shows a small RMSD of 0.548 Å for the backbone atoms. This conformational flexibility also explains why crystals of this complex could not be obtained. The quality of the final 20 lowest-energy conformers was verified using MOLMOL (15) for NOE violations and PROCHECK (16) for Ramachandran statistics. Data on the 20 lowest-energy conformers have been deposited in the protein data bank.

Data were analyzed and plotted using CARA (11). CSP analysis was done according to published protocols (17), and CSPs were plotted onto the structure by PyMOL (http://www.pymol.org/).

**Generation of an avid $UbV^{W\_dim}$ trap for cryo EM.** Expanding on our coexpression strategy for generation of a controlled dimeric $UbV^{W\_dim}$ for structural analysis by NMR, we similarly developed this controlled dimer with N-terminal and C-terminal extensions as outlined in Fig. 4f. The wildtype protomer was cloned by Gibson Assembly to a pRSF vector containing an N-terminal, TEV-cleavable $His_6$ tag, residues 770-792 of S. cerevisiae Hsl1 (KEN box), $UbV^W$, and residues 197-222 of H. sapiens UBE2S (C-terminal tail):
GSSGVSTNKENEGPEYPTKIEKNQFNMQIFVDTVQWKTITLEVEPSDTIENVKAKIQDKE
GIPPDQQRLIFAGKQLEDGRTLSDYNIQKESALILLLTLRKKHAGERDKKLAAKKKTDKK
RALRRL.
The mutant protomer was cloned by Gibson Assembly to a pGEX vector containing an N-terminal, TEV-cleavable GST tag, FLAG tag used for the APC/C capture step, linker-flanked residues 820-842 of S. cerevisiae Hsl1 (Dbox), $UbV^R$, linker, $UbV^W$ with I44N mutation as such:
GSDYKDDDDKGSTKIEKYLEEQKPKRAALSDITNSFNKMNSGMQILVKTPRGKTITLEVE
PSDTIENVKAKIQDKEGIPPDQQILFFAVKRLEDGRTLSDYNIQKKSSLLLAMRVPGKMKS
GGASSGSMQIFVDTVQWKTITLEVEPSDTIENVKAKIQDKEGIPPDQQRLNFAGKQLEDG
RTLSDYNIQKESALILLLTLR.

**Purification and cryo EM of $APC/C^{CDH1}$ with $UbV^{W\_dim}$ trap.** For the complex representing $UbV^{W\_dim}$ association with APC2 WHB, APC/C was expressed with HRV14 3C protease-cleavable tags, a Twin-Strep tag at the N-terminus of APC2 and a GST-tag at the N-terminus of APC16, and the complex with CDH1 was purified as previously described (3). $APC/C^{CDH1}$ was incubated with the avid $UbV^{W\_dim}$ trap, Anti-FLAG affinity gel (Genscript), and HRV14 3C protease for 1 hour. The resin was thoroughly washed in microspin columns, and complexes were eluted with antigenic peptide.

For cryo electron microscopic studies of APC/C complexes, 125 µg of purified APC/C-$UbV^{W\_dim}$ trap was loaded onto a 10%–40% glycerol gradient containing 50 mM HEPES pH 8.0, 200 mM NaCl, 2 mM MgCl2. For particle fixation by GraFix (18), the gradient also contained 0.025% and 0.1% glutaraldehyde in the lighter and denser glycerol solution, respectively, creating an additional glutaraldehyde gradient from top to bottom (0.025–0.1%). Centrifugation was performed at 34,000 rpm in a SW55TI rotor (Beckman) for 15 hr at 4°C. For cryo EM the fractions containing APC/C were subjected to a buffer exchange procedure using Zeba spin columns (Pierce) to remove the sugar prior to EM grid preparation. APC/C particles were allowed to adsorb on a thin film of carbon for 5 min, transferred onto a cryo EM grid (Quantifoil R2/1, Jena) and then plunged into liquid ethane under controlled environmental conditions of 4 °C and 100% humidity in a vitrification device (Vitrobot Mark IV, FEI Company, Eindhoven).

Micrographs were recorded on a K2 direct detector (operated in electron counting mode) under liquid-nitrogen conditions with a Titan Krios electron microscope (FEI, Eindhoven) equipped with a GIF quantum energy filter (20 $e^{-}V$) (Gatan). SerialEM (19) package was used for automated data acquisition with an electron dose of ~46 electrons (collected in 40 fractions) per $Å^2$, −0.7 to −3.3 µm defocus and a nominal magnification of 105,000X, resulting in a final pixel size of ~1.34 Å. The dose-fractionated movies were gain normalized, aligned and dose-weighted using MotionCor algorithm implemented in relion-3.0(20). CTFFIND4 (21) was used to determine the defocus values and particles were picked by Gautomatch (22) using a template derived from a smaller dataset. Particles were extracted and subjected to 2D classification in Relion to remove any contaminations. Particles were then taken through 3D classification and the class with greatest occupancy for UbV trap was auto-refined in Relion. This process was repeated iteratively yielding a final reconstruction resolution of 6.6 Å, which was estimated by applying a soft mask around the protein density and using the gold standard Fourier shell correction (FSC) = 0.143 criterion, as implemented in the RELION post-processing routine. Coordinates for APC/C with APC2 and APC11 deleted were placed as a single unit into the cryo EM maps alongside APC2 N-549 (PDB 4UI9,(4)) and composite CDH1-Dbox-KEN box (PDB 5KHR,(23)) using Chimera (24). APC2 residues 560-730 with APC11 residues 1-16 (PDB 4UI9) were fit in separately using Chimera. Extra density was modeled manually with considerations for distance restraints created by genetic linkages and known associations of domains with APC/C$^{CDH1}$ using the UbV$^{W}$-$^{dim}$ structure obtained in this study and UbV$^{R}$~APC11 (PDB 5JG6,(25)).

**Assaying APC/C substrate degradation in Xenopus egg extracts.** Interphase Xenopus egg extract was prepared as described (5, 26). Cyclin B1 Δ90 was added at 300nM and incubated for 120 minutes at room-temperature. Ubiquitin or UbV$^{W}$ were added at 10μM and incubated for 10 min. 110 nM full-length Cyclin B1/CDK1 or Securin was then added. To monitor substrate degradation, samples were diluted in SDS-PAGE sample buffer at 0, 30, and 60 minutes post substrate addition and processed for SDS-PAGE and immunoblotting using antibodies raised against Human Cyclin B1, Securin, APC3, and Smc3.

**X-ray crystallography.** The APC2 WHB + UbV$^{W}$ complex for crystallization was purified by co-mixing purified components separated from tags, and subsequent treatment to SEC in a final buffer of 20 mM Tris pH 7.6, 150 mM NaCl, 1 mM TCEP and frozen at 56mg/mL. 200 nL of this sample was mixed 1:1 (v:v) for crystallization by the hanging-drop vapor diffusion method. The reservoir solution contained 0.1 M Sodium Acetate pH 4.6, 0.02 M Calcium Chloride, 30% MPD, and 0.1 M Potassium Sodium Tartate Tetrahydrate. Despite the presence of APC2 WHB in the crystallization drop, crystals only contained UbV$^{W}$; no electron density was observed for APC2 WHB. The crystal was flash-frozen with liquid nitrogen in mother liquor prior to data collection at the SER-CAT Sector 22 ID beamline.

The APC2 WHB + Ubiquitin complex for crystallization was created by co-mixing purified components; APC2 WHB was separated from its tag, and each protein was then individually treated to SEC in 50 mM Tris pH 7.6, 200 mM NaCl, 1 mM DTT. 2.5 mM of pure APC2 WHB was mixed with 2.08 mM Ubiquitin on ice >30min prior to crystallization. 2μL of this sample was mixed 1:1 (v:v) for crystallization by the hanging-drop vapor diffusion method. The reservoir solution contained 0.1 M HEPES pH 7.7, and 27.5% PEG 3350. For cryoprotection, crystals were transferred to a neighboring drop containing reservoir solution supplemented with 30% PEG 400 prior to flash-freezing in liquid nitrogen. Data were collected at the NE-CAT Sector 24 ID C beamline.

Diffraction data were processed with HKL2000 for UbV$^{W}$ and XDS for the APC2 WHB + Ubiquitin complex (27);(28). The UbV$^{W}$ structure was determined by molecular replacement (MR) using Phaser (29) with a Ub search model, Protein Data Bank (PDB) ID code 4S1Z. The APC2 WHB + UbV$^{W}$ complex structure was determined by MR using Phaser with APC2 WHB from PDB ID code 4YII, chain C, and Ub (1-70) from PDB ID code 4S1Z as search models(3),(30). Model building and refinement were performed using Coot, Refmac5, and

Phenix (31), (32),(33). Data collection and refinement statistics are provided in Supplemental Table S1.

**Bio-layer interferometry (BLI).** Concentrated analyte and ligand proteins were diluted into BLI reaction buffer (25 mM HEPES pH 8, 200 mM NaCl, 0.01% Tween20). BLI experiments were performed on an Octet RED96 system (ForteBio) using anti-GST antibody biosensors for GST-tagged APC2 WHB, and untagged $UbV^{W}$ or $UbV^{W\_dim}$ at 25°C. A 9-fold dilution series ($UbV^{W}$) or 8-fold dilution series ($UbV^{W\_dim}$) covering a wide concentration range was applied for n≥3 experiments. Sensorgram raw data was processed and extracted by Octet Analysis 9.0 software.

**Size-exclusion chromatography-multi angle light scattering.** Purified proteins used in light scattering experiments were buffer exchanged identically to 20 mM HEPES pH 8, 150 mM NaCl, 1 mM TCEP buffer. The molecular masses were determined using a WTC-030S5 column (Wyatt Technology Corporation, Santa Barbara, CA, U.S.A.) attached to a miniDAWN® TREOS® static light scattering detector and an Optilab T-rEX refractive index detector (both Wyatt Technology) equilibrated to 20 mM HEPES pH 8, 150 mM NaCl, 1 mM TCEP at 0.5 mL/min flowrate. Data analysis was performed with ASTRA Softare (Version 6.1.7.17) and retraced manually.

**Analytical ultracentrifugation (AUC).** Purified proteins used in analytical ultracentrifugation experiments were buffer exchanged identically to 20 mM HEPES pH 8, 150 mM NaCl, 1 mM TCEP buffer. Sedimentation velocity experiments were performed on an Optima XL-I analytical centrifuge (Beckman Inc., Indianapolis, IN, U.S.A.) using an A 60-Ti rotor and double-sector 12 mm centrepieces. Buffer densities were measured using a DMA 5000 densitometer (Anton Paar, Graz, Austria). Protein concentration distribution was monitored at 280 nm, using 50,000 r.p.m. Time-derivative analysis was computed using the SEDFIT software package (34), resulting in a c(s*) distribution and an estimate for the molecular weight (from sedimentation coefficient and the diffusion coefficient, inferred from the peak width).

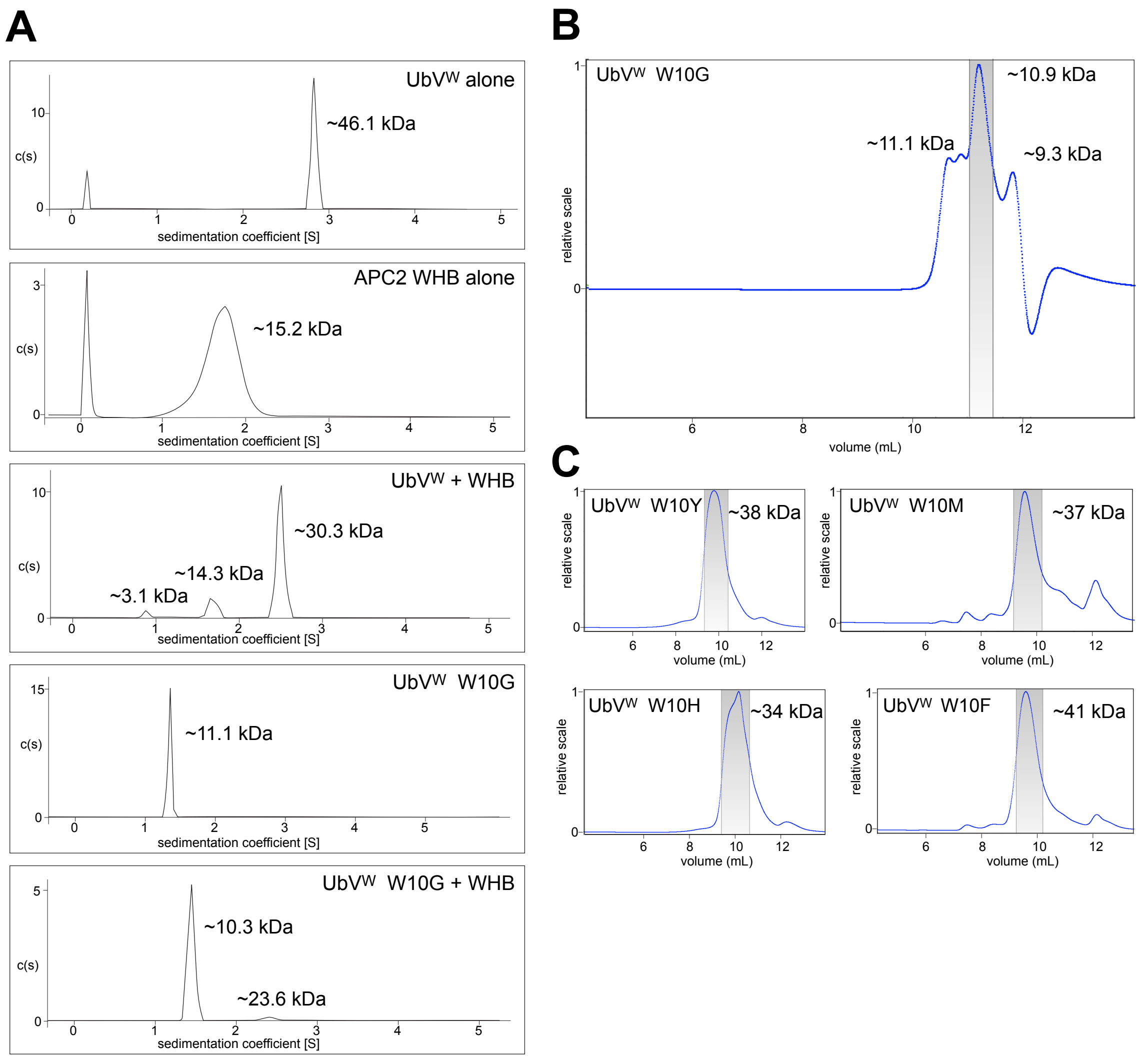

**Fig. S1. Biophysical characterization of UbV mutants.** *(A)* Representative analytical ultracentrifugation data for UbV variants, APC2 WHB domain, or complexes. *(B)* Size-exclusion chromatography-multi-angle light scattering of UbV$^{W}$ Trp10 reversion to Glycine (as in Ubiquitin) yields an ~11kDa species. *(C)* Size-exclusion chromatography-multi-angle light scattering for Tyr, Met, His, or Phe substitutions for UbV$^{W}$ Trp10 yield species consistent with oligomerization.

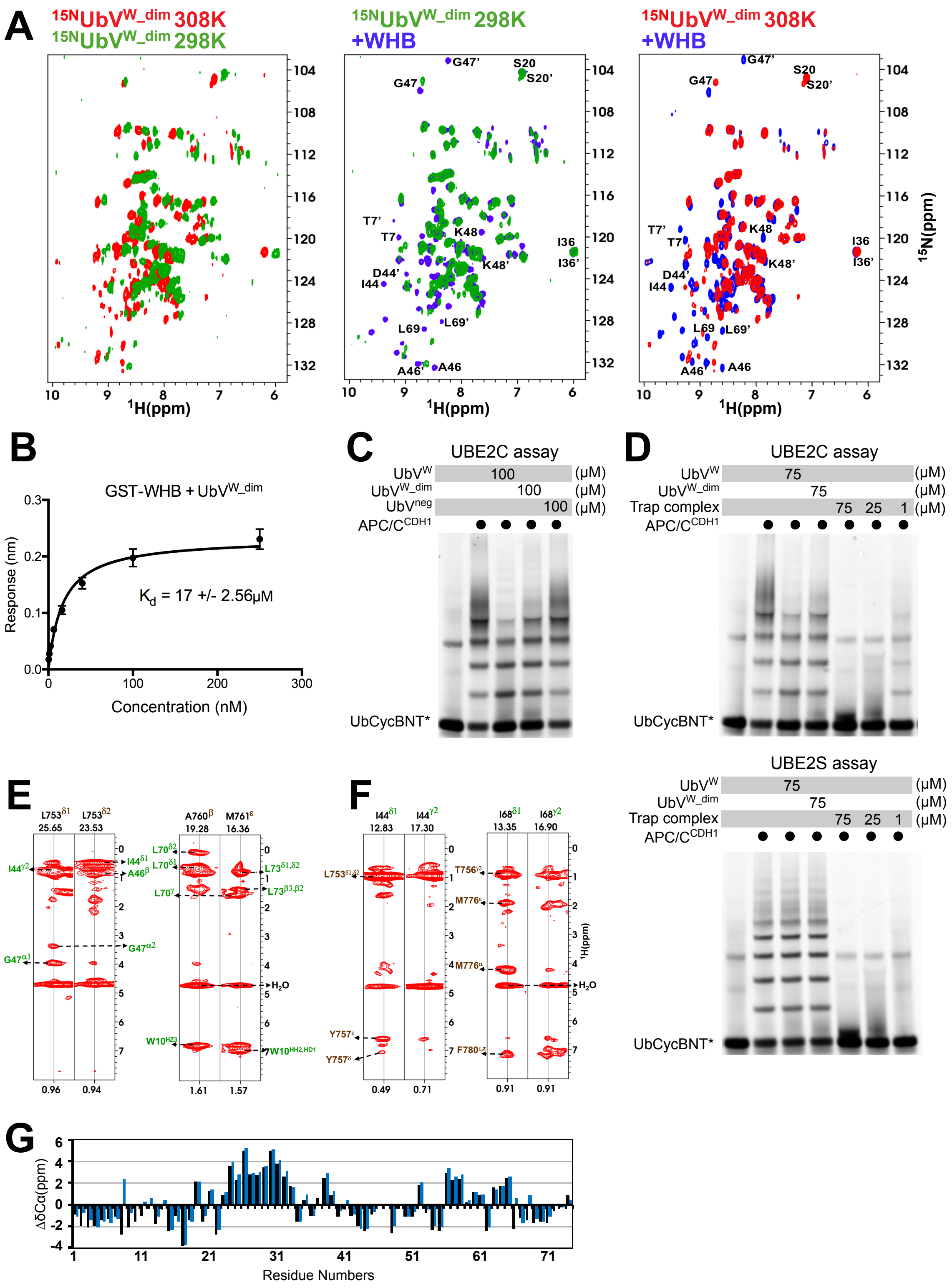

**Fig. S2. Enzymatic and NMR characterization of UbV$^{W_dim}$.** *(A)* Left, overlaid 2D [$^{15}$N, $^{1}$H] TROSY spectra of UbV$^{W_dim}$ at 298K or 308K sampling temperatures; several resonances become sharper at 308K. Middle, overlaid 2D [$^{15}$N, $^{1}$H] TROSY spectra of UbV$^{W_dim}$ either in apo form (green) or in the presence of WHB (blue) at 1:2 molar ratio, measured at 298K. Peaks from protomer B (I44D) labeled with apostrophe. Right, overlaid 2D [$^{15}$N, $^{1}$H] TROSY spectra of

$UbV^{W\_dim}$ either in apo form (red) or in the presence of WHB (blue) at 1:2 molar ratio, measured at 308K. Peaks from protomer B (I44D) labeled with apostrophe. Several apo resonances of $UbV^{I44D}$ were missing (eg. G47‘) even at 308K due to exchange broadening, and binding of WHB stabilized the dynamics enabling the resonances to be seen. *(B)* For comparison to APC2 WHB domain binding to $UbV^{W}$ as shown in Fig.2d, representative curve fit for binding measured by bio-layer interferometry (BLI) with soluble $UbV^{W\_dim}$ and immobilized GST-WHB domain. SEM, n ≥ 3. *(C)* Ubiquitylation of UbCycBNT* by the APC/C is monitored in the presence of 75μM $UbV^{W}$, $UbV^{W\_dim}$, or at the indicated concentrations of the trap complex in UBE2C (top) or UBE2S (bottom) reactions. The trap complex, shown schematically in Figure 4F, comprised of substrate D-box, $UbV^{R}$, $UbV^{W}$ protomer B and $UbV^{W}$ protomer A with KEN box and UBE2S tail, inhibits both E2 reactions at 1μM. *(D)* Ubiquitylation of UbCycBNT* by the APC/C is monitored in the presence of 75μM $UbV^{W}$, $UbV^{W\_dim}$, or at the indicated concentrations of the trap complex (as in Fig. 4) in UBE2C (top) or UBE2S (bottom) reactions. The trap complex, comprised of substrate D-box, $UbV^{R}$, $UbV^{W}$ protomer B and $UbV^{W}$ protomer A with KEN box and UBE2S tail, inhibits both E2 reactions at 1μM. *(E)* Intermolecular NOEs observed between $^{15N}$WHB (brown) and unlabeled $UbV^{W\_dim}$ (green). NOEs are observed for the indicated resonances of WHB residues labeled at the top. *(F)* Intermolecular NOEs observed between $^{15N}UbV^{W\_dim}$ (green) and unlabeled WHB (brown). NOEs are observed for the indicated resonances of $UbV^{W\_dim}$ residues labeled at the top. *(G)* Cα deviation plot versus residue number shown for $UbV^{W}$-APC2 WHB (black) or Ubiquitin monomer (blue). The linker region between β1 and β2 strands in Ubiquitin (residues 8-14) have changed to negative values in the complex, suggesting a long strand or domain-swapped conformation of $UbV^{W}$.

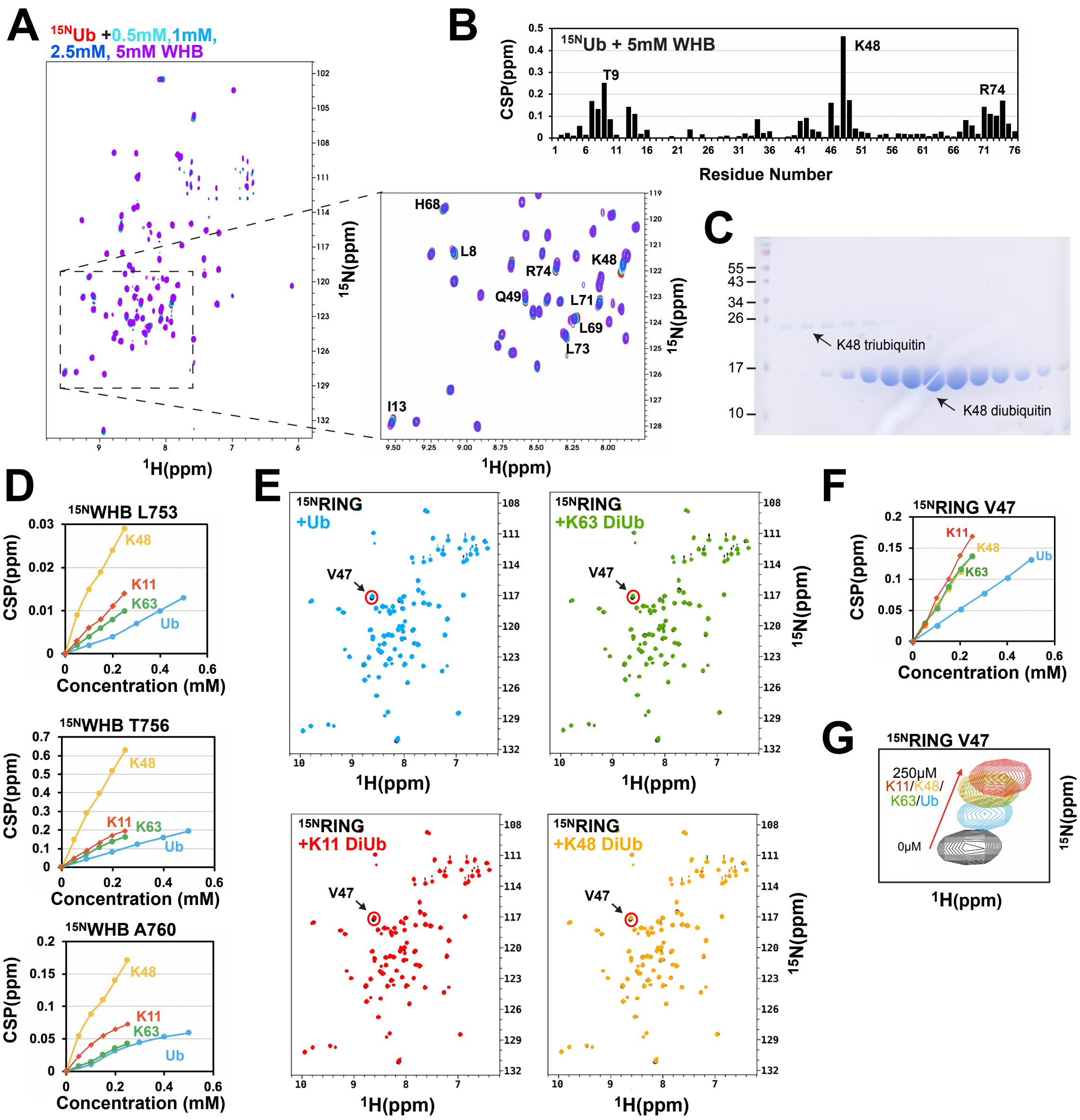

**Fig. S3. Analysis of Ubiquitin association with APC2 and APC11.** *A)* Overlaid 2D [$^{15}$N, $^{1}$H] TROSY spectra of 100µM $^{15N}$Ubiquitin apo (red) or with the indicated concentrations of APC2 WHB domain. Isolated APC2 WHB causes chemical shift perturbation of several Ub residues. A subset of residues are labeled in the zoomed subregion. *(B)* CSP analysis for 100µM $^{15N}$Ub in complex with 5mM isolated APC2 WHB compared to apo shows perturbation of residues contributing to the hydrophobic binding surface on Ubiquitin. *(C)* Coommassie-stained 16.5% SDS-PAGE analysis of diubiquitin purification illustrates typical diubiquitin purity for NMR titrations. Lanes 2-15 contain concurrent fractions from SEC. (*D*) As in Fig. 5c, CSP values at the indicated residues of $^{15N}$WHB are plotted for titrations of K48-, K11-, or K63- diubiquitin and purified Ubiquitin. *(E)* Overlaid [$^{15}$N, $^{1}$H] HSQC spectra of $^{15}$N-labeled APC11 RING domain alone (200µM, black) or: with Ubiquitin (1:2.5, light blue), with K63 diubiquitin (1:1.25, green), with K11 diubiquitin (1:1.25, red), or with K48 diubiquitin (1:1.25, orange). Val47 is marked as a dominant chemical shift and was previously published for Ubiquitin. *(F)* CSP values for Val47 of $^{15N}$RING domain are plotted for titrations of K48-, K11-, or K63- diubiquitin and purified

Ubiquitin. No diubiquitin preference is observed. *(G)* Closeup analysis of spectral point for residue Val47 at the Ubiquitin-APC11 RING interface at a single titration concentration with indicated titrants. No diubiquitin preference is observed.

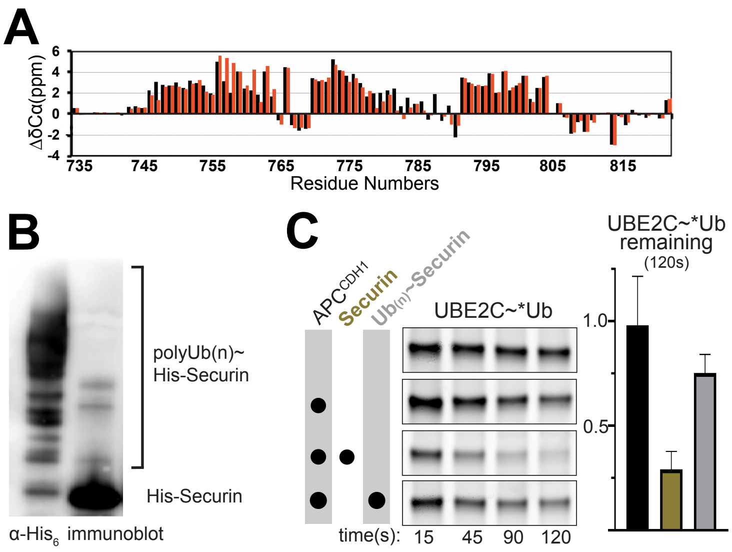

**Fig. S4. Validation of Ubiquitin association with APC2 WHB.** *(A)* Cα deviation plot versus residue number shown for APC2 WHB apo-(black) or bound to $UbV^{W}$ (red). Deviations in the C-terminal region of α1 and the loop connecting α2 and α3 suggest possible conformational changes in these regions due to association of $UbV^{W}$, and potentially Ub. *(B)* Normalized loads for polyubiquitylated Securin and Securin visualized by western blot for His tag. *(C)* Left, timecourse of pulse-chase assays monitoring UBE2C discharge of fluorescent Ub (*Ub) in the presence of $APC/C^{CDH1}$ and either Securin (gold) or polyubiquitylated Securin (gray). Right, quantitation of the 120s timepoint of the indicated remaining UBE2C~Ub.

**Table S1. Crystallogaphy Data Collection and Refinement Statistics.**

| | $UbV^{W}$ | Ub-APC2 WHB |
|---|---|---|
| Data collection | | |
| Space group | $P4_1$ | $P2_12_12_1$ |
| Cell dimensions | | |
| *a*, *b*, *c* (Å) | 62.36, 62.36, 168.12 | 58.93, 63.37, 99.88 |
| α, β, γ (°) | 90.0, 90.0, 90.0 | 90.0, 90.0, 90.0 |
| Resolution (Å) | 30.0-2.80 (2.85-2.80) | 53.55-2.20 (2.27-2.20) |
| $R_{\text{pim}}$ (%) | 5.1 (16.9) | 3.6 (30.3) |
| $R_{sym}$ (%) | 8.4 (23.7) | 6.9 (58.3) |
| $I/\sigma I$ | 14.8 (2.0) | 12.0 (2.3) |
| Completeness (%) | 96.1 (67.0) | 96.5 (99.0) |
| Redundancy | 3.5 (2.1) | 4.4 (4.4) |
| Refinement | | |
| Resolution (Å) | 29.6-2.80 (2.98-2.80) | 43.55-2.20 (2.26-2.20) |
| No. reflections | 15,083 (1,933) | 17,815 (1,335) |
| $R_{\text{work}}/R_{\text{free}}$* | 21.34/26.54 (30.9/38.3) | 22.05/26.62 (33.31/30.80) |
| No. molecules in ASU | 8 | 4 (2 Ub, 2 APC2 WHB) |
| No. atoms | | |
| Protein | 4,490 | 2,403 |
| Water | 0 | 116 |
| B factors | | |
| Protein | 66.20 | 49.31 |
| Water | N/A | 50.70 |
| Rmsd | | |
| Bond lengths (Å) | 0.009 | 0.010 |
| Bond angles (°) | 1.24 | 1.34 |

- $R_{\text{pim}}$, redundancy-independent merging R factor; ASU, asymmetric units. Data for outer shell shown in parentheses.
- *$R_{\text{free}}$ test set size 5.2 %

**Table S2. NMR Structural Statistics for $UbV^{W\_dim}$.**

| Parameters | WHB | $UbVw^{I44}$ | $UbVw^{D44'}$ |
|---|---|---|---|
| **Distance constraints** | | | |
| Total number of NOEs | 1062 | 914 | 892 |
| Intramolecular | 333 | 237 | 232 |
| Short range (\|i-j\| ≤ 1) | 309 | 276 | 277 |
| Medium-range (1 ≤ \|i-j\| ≤ 5) | 226 | 138 | 131 |
| Long-range (\|i-j\| ≥ 5) | 194 | 263 | 252 |
| Intermolecular NOEs | | | |
| $UbV^{I44}$ | | 55 | |
| $UbV^{D144}$ | | | 2 |
| Hydrogen bonds | 172 | 134 | 128 |
| Total dihedral angle constraints | | | |
| Phi (ϕ) | 61 | 59 | 59 |
| Psi (ψ) | 61 | 59 | 59 |
| Violations | | | |
| Upper Distance (ave) (Å) | 0.014 ± | | |
| Lower Distance (ave) (Å) | 0.08 | | |
| Angle (ave) (°) | 0.005 ± | | |
| | 0.01 | | |
| **Average pairwise r.m.s. deviation** | 0.589 ± | | |
| **(Å)**[a] | 0.10 | | 0.55 ± |
| Backbone | | | 0.11[b] |
| Heavy atom | | | 1.08 ± 0.13 |
| | 0.64 ± 0.13 | | |
| **Ramachandran statistics**[a] | 1.01 ± 0.12 | | |
| Residues in most favorable region (%) | | | |
| Residues in additional favorable | 82.7 % | | |
| region (%) | 14.9 % | | |
| Residues in generously allowed | 2.1 % | | |
| region (%) | 0.4 % | | |
| Residues in disallowed region (%) | | | |

[a]Heavy atom and backbone RMSD is calculated for residues 1'-9',10-72 of $UbV^{I44D}$ and 742-818 WHB. [b]Heavy atom and backbone RMSD is calculated for residues 1-9, 10'-72' of the second $UbV^{I44D}$ (D44') which is not involved in binding with WHB.
